# Cell-type-specific cortical feedback coordinates hierarchical credit assignment

**DOI:** 10.64898/2026.06.16.732595

**Authors:** Will Greedy, Heng Wei Zhu, Alexia Duriez, Joseph Pemberton, Patrick T. McCarthy, Kevin Kermani Nejad, Rui Ponte Costa

## Abstract

Learning is thought to arise from synaptic modifications across brain-wide circuits, yet how these networks coordinate plasticity to support complex behaviour is not known. Inspired by deep learning, we introduce a theory of hierarchical credit assignment in which pathway-specific cortical feedback drives dendrite-dependent burst plasticity across the cortex. We show that this principle enables learning in dynamic settings, complex visual recognition, and goal-directed tasks, mechanistically linking credit assignment to cell-type-specific regulation of dendritic excitation–inhibition balance. Our framework provides a unified account of diverse experimental phenomena – including cell-type-specific modulation of synaptic plasticity, learning-dependent changes in interneurons, and neuron-specific dendritic error signals. Furthermore, the theory predicts that interneurons constrain the dimensionality of error feedback, offering a functional rationale for cortex-wide gradients in interneuron density. Together, these findings reveal how distinct cortical cell types cooperatively coordinate learning, bridging the gap between synaptic plasticity, circuit-level computation, and behaviour.

## Introduction

To learn effectively, the brain must translate behavioural outcomes into synapse-specific changes across distributed neural circuits. This is a problem shared by artificial and biological hierarchical networks: the contribution of a local connection to behaviour or task performance only becomes apparent after activity has propagated through many downstream areas ^1,2^. In artificial neural networks, algorithms such as backpropagation solve this problem by explicitly computing and propagating credit signals backwards across many areas of processing ^3^. The brain must meet an analogous computational demand under biological constraints: synapses are modified using local information, while information about behavioural outcomes is available only indirectly through feedback pathways from downstream activity. This poses a central problem for neuroscience: how can biological circuits generate synapse-specific learning signals across complex distributed cortical hierarchies ^4^?

The cortex contains several features that support such a biological solution. Pyramidal neurons segregate feed-forward and feedback inputs across basal and apical dendritic compartments, allowing sensory drive and feedback-related signals to interact within single neurons ^5–8^. Apical input can regulate nonlinear dendritic events and somatic bursting ^9^, while bursting can determine the sign of long-term synaptic plasticity ^10^. These observations have motivated models in which apical dendrites encode credit signals through mismatches between local interneuron activity and downstream feedback ^5^, or through dendrite-mediated bursting ^11^. However, these existing approaches face critical limitations: they either fail to propagate error signals effectively through hierarchical cortical networks or rely on multi-phase learning that requires separation of inference from synaptic plasticity ^12^.

Here we propose that differential cortical feedback provides a circuit principle for assigning credit across cortical hierarchies. Inspired by credit-assignment approaches in AI, we formalise this principle in Bursting Cortico-Cortical Networks (BurstCCN), in which two dendrite-targeting feedback pathways jointly regulate somatic bursting to drive dendrite-dependent synaptic plasticity ^5,11,12^. By combining cell-type-specific feedback, burst multiplexing, short-term plasticity and burst-dependent plasticity, we show that BurstCCN enables online credit assignment across hierarchical cortical networks. In doing so, it brings together disparate observations spanning synaptic plasticity, interneuron function, and dendritic computation, while linking the computational power of deep learning to concrete features of cortical microcircuits.

## Results

### Differential cortical feedback enables burst-mediated credit assignment

To improve behavioural performance, credit assignment must ultimately drive local changes in synaptic weights. Experimental evidence indicates that postsynaptic activity determines the sign of long-term plasticity, with strong bursting activity inducing long-term potentiation (LTP) and weaker activity inducing long-term depression (LTD) ^10,13–17^. Inspired by these experimental observations and previous models ^11,18^, we implemented a burst-dependent plasticity rule that controls the sign of synaptic change. This rule considers two types of spiking events: *single-spike* and *burst* events. The fraction of events that are burst events is referred to as the *burst probability* ^18^. The burst-dependent plasticity rule is given by

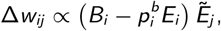

where *E*_*i*_ and *B*_*i*_ are binary indicators of postsynaptic events and burst events, respectively: *E*_*i*_ = 1 for either single-spike or burst events, *B*_*i*_ = 1 for burst events only, and both are 0 otherwise. Here, 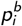 is a constant that we interpret as a baseline burst probability, and 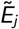 is a presynaptic event trace (see Methods). The sign of plasticity is therefore determined by the type of postsynaptic spiking event: burst events contribute a positive factor, 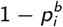, whereas isolated single-spike events contribute a negative factor, 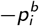. When averaged over time, this postsynaptic contribution is proportional to 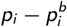, where *p*_*i*_ is the burst probability (Fig. 1; cf. Eq. 5). Therefore, 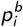 acts as a threshold that separates potentiation from depression, in line with threshold-dependent mechanisms commonly employed in synaptic plasticity models ^17,19,20^.

**Figure 1.**
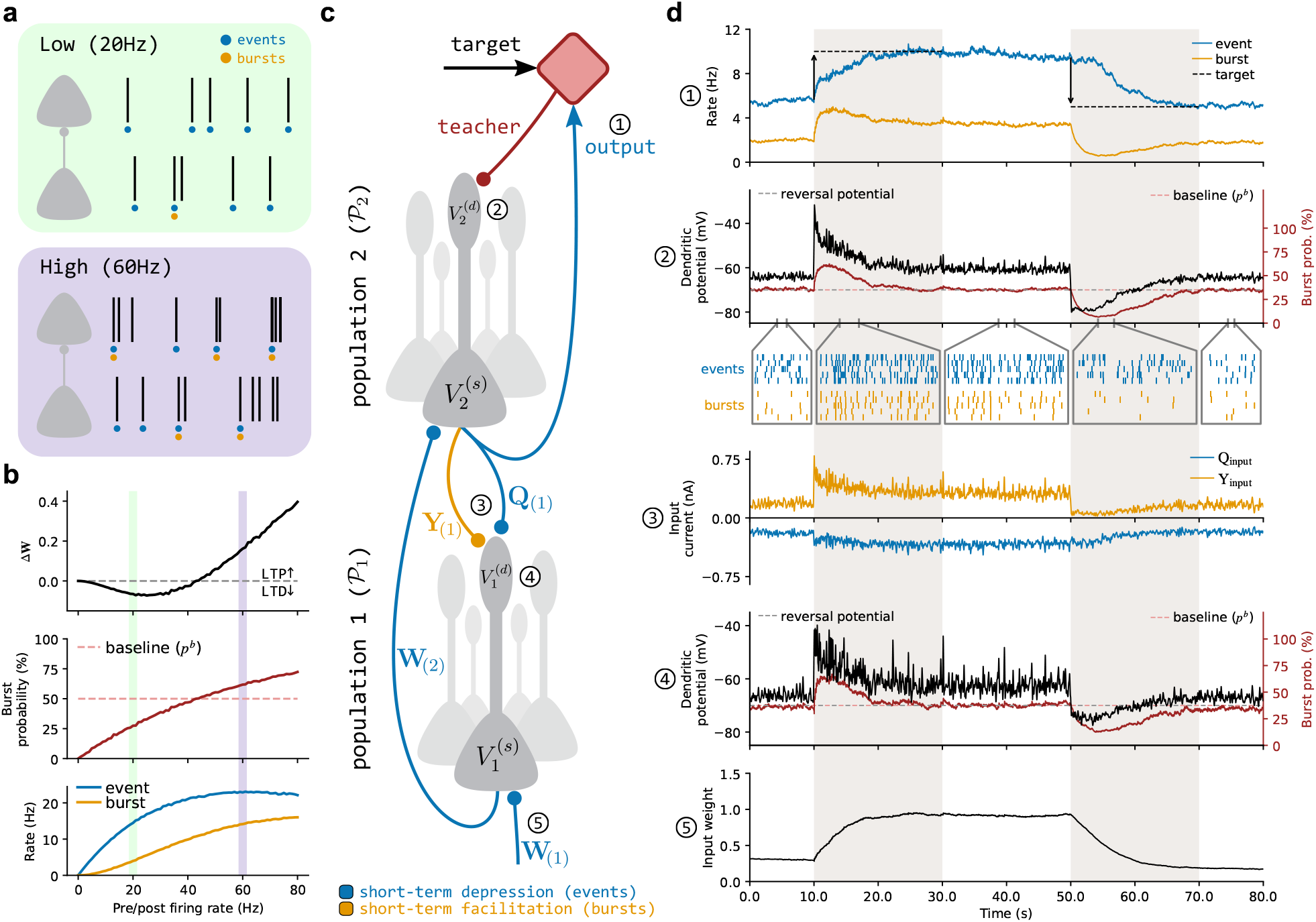
Differential cortical feedback enables burst-dependent synaptic plasticity. **a**, Illustration of two Poisson spiking neurons connected by a single synapse. Each spiking event is indicated by a blue dot, with the subset of burst events marked by orange dots. Low firing rates (top) produce fewer burst events than high firing rates (bottom). **b**, Burst-dependent plasticity as a function of firing rate. Top: As the firing rates of both neurons increase from low to high, the sign of synaptic plasticity switches from LTD to LTP. Middle: This switch occurs when the postsynaptic neuron’s burst probability (*p*, red line) crosses the baseline burst probability (*p*_*b*_, dashed red line). Bottom: The increase in burst probability increases the likelihood of burst events (orange) relative to all spiking events (blue). **c**, Two-population spiking network used for the stationary target-prediction task. Blue connections denote event-decoding feedback pathways mediated by short-term depression, and orange connections denote burst-decoding feedback pathways mediated by short-term facilitation. **d**, Network state variables over time during the task. Grey areas denote periods in which the output population is given a target event rate: a high target rate (10 Hz, 10–30 s) followed by a low target rate (5 Hz, 50–70 s). During these periods, (1) the mismatch between the target rate and the event rate of the second population, *P*_2_, generates a teaching signal. This (2) shifts burst probability away from baseline and (3) changes the burst-feedback input, **Y**_input_, received by the apical dendrites of the first population, *P*_1_. (4) This feedback changes dendritic potential and bursting in *P*_1_, which (5) induces long-term plasticity in the input weights, **W**_(1)_. Plasticity continues until the event rate of *P*_2_ matches the target rate.

To illustrate the behaviour of this burst-dependent plasticity rule, we first considered a minimal setting consisting of two Poisson spiking neurons connected by a single synapse. We compared low (20 Hz) and high (60 Hz) firing regimes for both pre- and postsynaptic neurons (Fig. 1a). In the low-rate regime, single-spike events were more frequent, such that plasticity was dominated by LTD (Fig. 1b). As the firing rates of both cells increased from low to high, burst events became more frequent, increasing the burst probability. This increase in bursting drove a switch in net plasticity from LTD to LTP once the burst probability crossed its baseline value 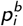.

For burst-dependent plasticity to support task-relevant learning, burst probability must carry information beyond overall firing activity and be shaped by credit-related feedback from downstream areas. Experimental evidence indicates that bursting is a distinct firing mode that can encode information independently of spike rate ^18,21^. Consistent with this, Naud and Sprekeler ^22^ proposed a burst-multiplexed code in which event rate reflects feedforward drive to basal dendrites, whereas burst probability encodes feedback drive to apical dendrites. This is supported by observations showing that cortical feedback inputs targeting apical dendrites modulate burst generation in pyramidal neurons ^9,23^. Event and burst signals can then be decoded through short-term plasticity, with short-term depression (STD) decoding events and short-term facilitation (STF) preferentially decoding bursts ^22^.

Building on these principles, we next illustrate how BurstCCN combines burst-dependent plasticity, burst multiplexing and apical feedback to support biologically plausible credit assignment in a two-population spiking network (Fig. 1c). This network was tasked with adjusting the event rate of the output population, *P*_2_, towards a target rate. The input population, *P*_1_, received constant basal input through weights **W**_(1)_, which drove its event rate. This activity was propagated to the basal dendrites of *P*_2_ through event-decoding STD synapses, **W**_(2)_. The *P*_2_ neurons received an apical teaching signal proportional to the mismatch between the target rate and their current event rate, which modulated their burst probability. To test whether this teaching signal could drive upstream plasticity, we fixed **W**_(2)_ and restricted plasticity to **W**_(1)_. The feedback is mediated by two apical-targeting pathways that decode distinct components: the **Y** pathway decodes bursts via STF, while the **Q** pathway decodes events via STD. The apical compartment therefore compares a burst-dependent feedback input (**Y**_input_) with a burst-independent event-based reference input (**Q**_input_). When the downstream population is at its baseline burst probability, the **Q** and **Y** feedback weights are configured such that these inputs cancel, producing no net apical drive and therefore no net plasticity. We refer to this as the **QY**-symmetric configuration (see Methods).

We simulated this two-population network using the spiking implementation of BurstCCN (see Methods) and examined the network dynamics as it learned to match sequential high (10 Hz) and low (5 Hz) target event rates (Fig. 1d). When a high target event rate was introduced, (1) the mismatch with the current event rate generated a positive teaching signal that (2) increased the burst probability of *P*_2_, increasing burst-sensitive **Y** feedback and creating an imbalance between the **Q** and **Y** inputs to the apical dendrites of *P*_1_. This imbalance (4) increased dendritic depolarisation and burst probability in *P*_1_ (*p > p*^*b*^), resulting in (5) potentiation of the feedforward weights, **W**_(1)_. This potentiation drove *P*_2_ to match the target output, reducing the teaching signal towards zero, allowing burst probability to return to baseline and stopping further plasticity. Importantly, the learned output remained stable once the teaching signal was removed. Introducing a lower target event rate engaged the same mechanism in the opposite direction, reducing *P*_1_ burst probability below baseline (*p < p*^*b*^) and inducing LTD at its basal synapses.

These results demonstrate the core mechanism behind BurstCCN: differential cortical feedback propagates downstream teaching signals to upstream dendrites, where deviations in burst probability drive local synaptic plasticity.

### Differential cortical feedback enables online hierarchical credit assignment

A key advantage of BurstCCN is that learning does not require temporally distinct inference and teaching phases. The event-sensitive **Q** pathway provides a continuous reference signal, while the burst-sensitive **Y** pathway simultaneously conveys teaching-related deviations from this reference, allowing each neuron to determine the sign of synaptic plasticity during stimulus presentation. To test this single-phase learning more directly, we trained the spiking BurstCCN on XOR, a nonlinear associative task that requires plasticity in intermediate populations (Fig. S1a–d). BurstCCN learned the task without separate inference and teaching phases, demonstrating that differential feedback can support single-phase learning of input-dependent associations in this biologically grounded spiking model.

We next asked whether the same principle supports learning in fully continuous-time settings, where both inputs and targets evolve dynamically. This is important for naturalistic learning, where sensory inputs and teaching signals are not presented in discrete trials but can change continuously over time. To demonstrate this behaviour, we used a continuous-time, rate-based implementation of BurstCCN (see Methods). We defined two nonlinear regression tasks in which, for each task, a predictor network was trained to match the output of a fixed target network with the same architecture (Fig. 2a), mimicking cortico-cortical learning in which one area learns to predict another ^25^.

**Figure 2.**
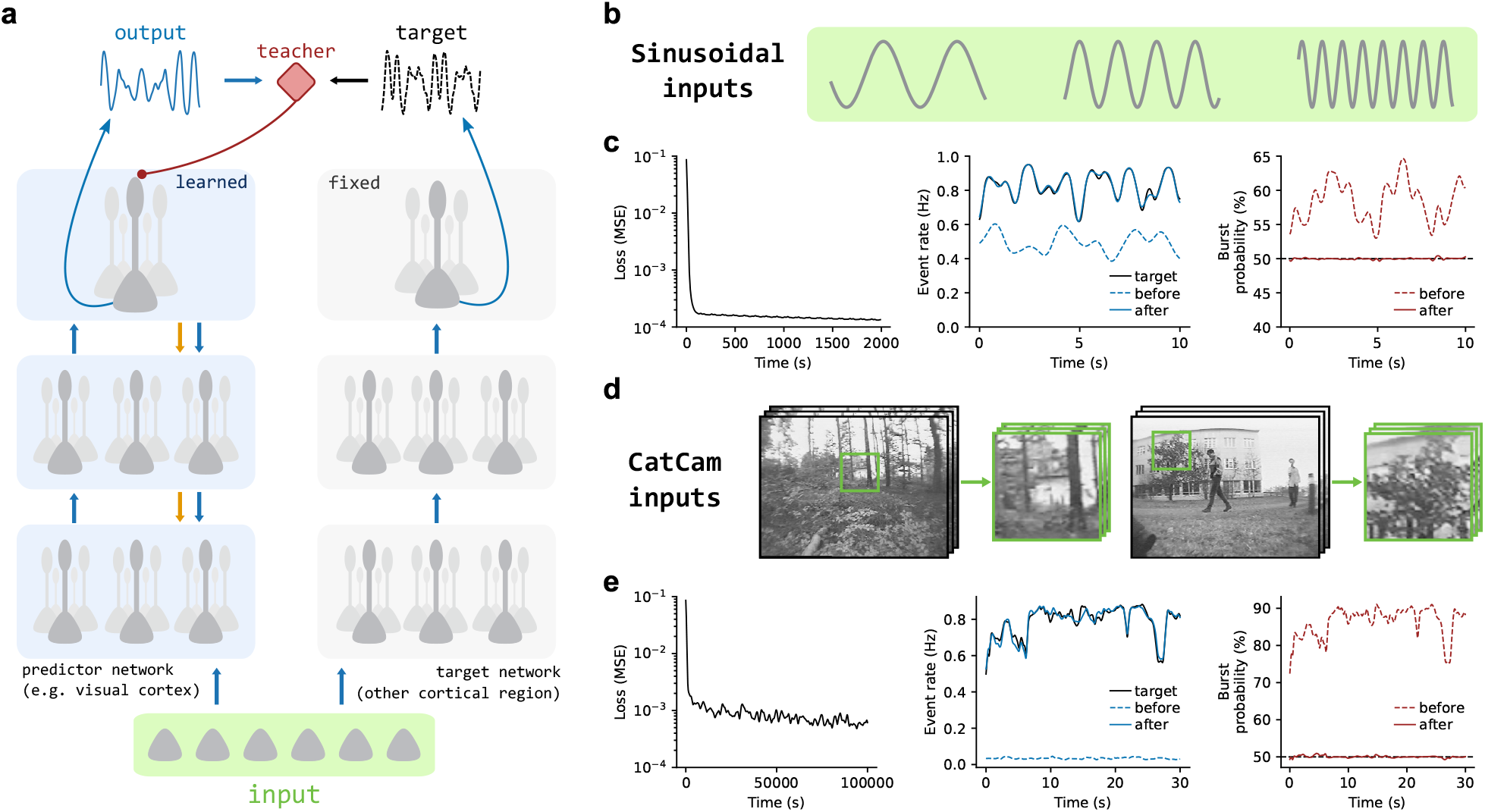
BurstCCN learns nonlinear dynamic regression tasks online. **a**, Illustration of the target signal generation process for each task. Copies of the same input are given to both the predictor network, which learns the task, and a separate randomly initialised fixed target network. **b**, Example inputs for the sinusoidal task. **c**, Performance and activity for the sinusoidal task. Left: mean-squared error across training. Middle: event-rate traces for the output neuron before and after training. Right: burst-probability traces for the same output neuron before and after training. The dashed black line indicates the baseline burst probability, *p*^*b*^ = 50%. **d**, Example inputs for the CatCam task ^24^. For each training video, a random cropped window was selected from the greyscale video (green box; see Methods). **e**, Performance and activity for the CatCam task, shown as in **c**. Activity is shown for one example output neuron; see Fig. S2 for all output neurons. In **c** and **e**, each line represents the mean, with standard error shown by the shaded area (*n* = 5). The images in (d) were obtained from publicly available datasets ^24^.

First, using sinusoidal inputs, a two-area predictor BurstCCN learned to reproduce the dynamically varying target output with minimal error despite its initially mismatched activity (Fig. 2b,c). We then tested the model with more naturalistic inputs using the CatCam dataset, which consists of outdoor naturalistic videos recorded from a cat’s perspective ^24^, converted into continuous-time inputs (Fig. 2d; see Methods). In a larger network with three areas and ten output neurons, each predictor output converged towards its corresponding target over training (Fig. 2e; Fig. S2). Together, these results show that the differential-feedback mechanism that enables online credit assignment in spiking networks also supports learning under more dynamic, naturalistic conditions.

### Dendritic-targeting plasticity enables feedback-pathway alignment

Having established the core mechanism in spiking and continuous-time settings, we used a discrete-time, rate-based implementation of BurstCCN for the remaining simulations, making it tractable to study deeper networks and more complex tasks. In all experiments presented thus far, feedback weights were fixed in a **QY**-symmetric configuration (see Methods), ensuring balanced event- and burst-feedback inputs in the absence of a teaching signal. This symmetry is important for enabling the propagation of learning signals across the network. However, it imposes a non-trivial biological constraint since establishing and maintaining this symmetry would suggest that synapses in one feedback pathway have access to non-local information relating to the strengths of synapses in the other, spatially distinct pathway.

In cortical circuits, balanced synaptic inputs are not imposed through fixed symmetry but instead develop through local, activity-dependent plasticity. Excitation–inhibition balance in both somatic and dendritic compartments is known to be dynamically maintained by inhibitory synaptic plasticity that depends only on locally available signals ^26,27^. Although the **Q** and **Y** pathways are not strictly excitatory or inhibitory, their alignment similarly balances net apical input in the absence of a teaching signal. Inspired by previous work ^28^, we therefore asked whether plasticity within the burst-feedback pathway alone is sufficient to establish this balance.

To test this, we restricted this new form of apical-targeting plasticity to the burst-feedback pathway, **Y**, while keeping the feedforward weights **W** and event-feedback weights **Q** fixed (Fig. 3a). We independently initialised **Q** and **Y** at random, so their apical inputs were initially mismatched and local **Y** plasticity was required to establish **QY**-symmetry. Plasticity of the **Y** weights followed a local apical learning rule, given by 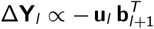, where **u**_*l*_ is the postsynaptic apical membrane potential and **b**_*l*+1_ is the presynaptic burst-rate feedback from the next area conveyed via the **Y** pathway. This rule acts to reduce net apical drive, restoring burst probability to its baseline level (Fig. 3b). Through plasticity, the burst-feedback pathway (**Y**) aligned with the fixed event-feedback pathway (**Q**), such that **Q** and **Y** inputs became balanced (Fig. 3c). Decomposing these inputs into their excitatory and inhibitory components further revealed that feedback plasticity produced a stimulus-specific excitation-inhibition balance of apical input across each input example (Fig. 3d), consistent with compartment-specific excitation–inhibition balance observed in cortical dendrites ^29–31^.

**Figure 3.**
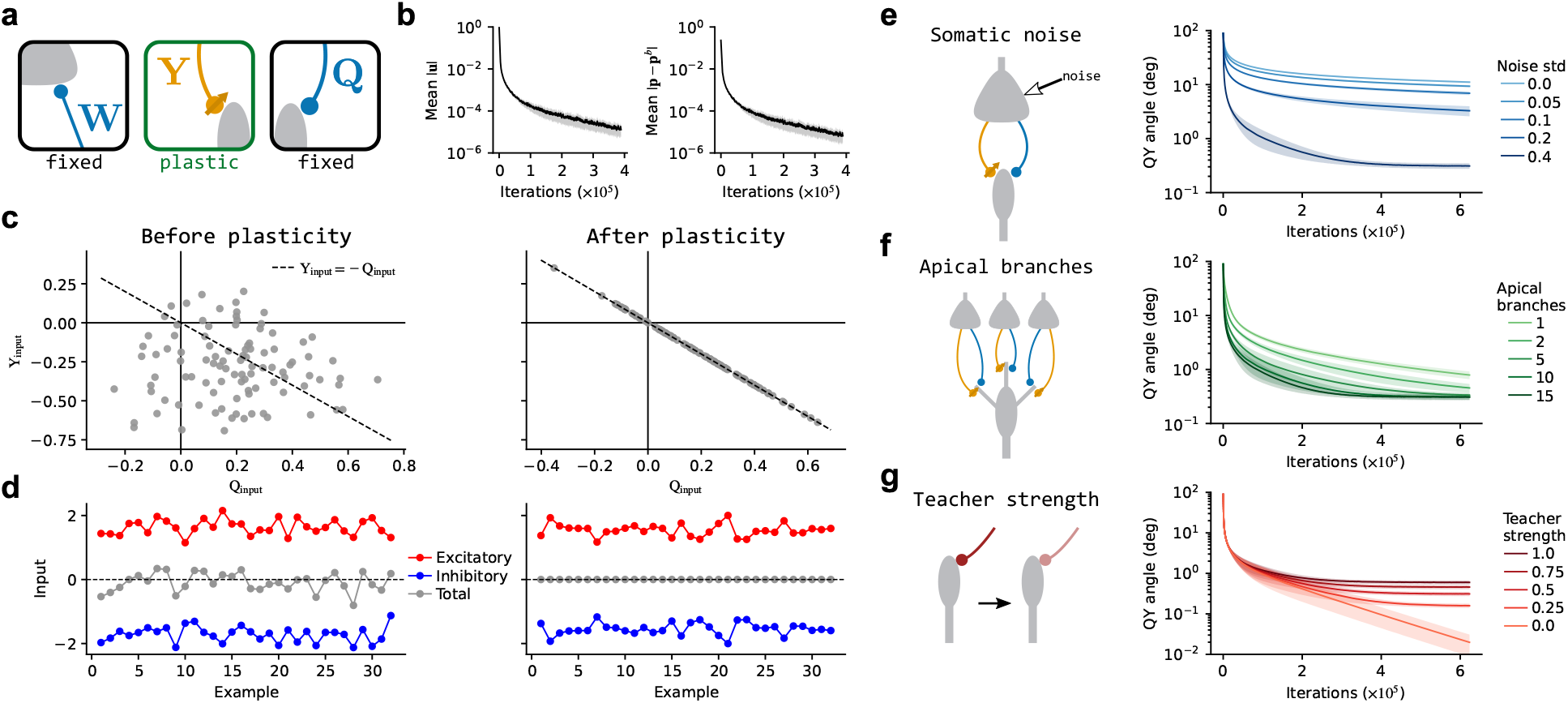
Dendritic-targeting plasticity enables feedback-pathway alignment. **a**, Burst-feedback plasticity configuration: burst-feedback weights **Y** are plastic, while feedforward weights **W** and event-feedback weights **Q** are fixed. **b**, Change from baseline in apical potential (left) and burst probability (right) during feedback-only plasticity. **c**, Total weighted inputs through **Q** (**Q**_input_) versus **Y** (**Y**_input_): before (left) and after (right) **Y** plasticity. The black dashed line denotes balance between **Q** and **Y** feedback inputs. **d**, Excitatory and inhibitory apical feedback inputs across feedforward input examples: before (left) and after (right) **Y** plasticity. **e**, Schematic of independent somatic noise injected into downstream neurons during **Y** plasticity (left). Angle between **Q** and **Y** weights during **Y** plasticity for different noise levels (right). **f**, Schematic of multiple apical branches receiving distinct **Q** and **Y** feedback inputs (left). Angle between **Q** and **Y** weights during **Y** plasticity for different branch numbers (right). **g**, Schematic of different teaching signal strengths to output apical dendrites (left). Angle between **Q** and **Y** weights during **Y** plasticity for different teaching signal strengths (right). Each line represents the mean, with standard error depicted by the shaded area (n = 5).

We next examined the key factors that determine the rate at which **QY**-alignment emerges. This alignment was quantified as the angle between the **Q** and **Y** weight matrices (see Methods), with smaller angles indicating greater functional alignment between the two pathways. First, we found that independent somatic noise in downstream neurons accelerated **QY**-alignment (Fig. 3e). Without noise, correlated activity within a downstream area allows burst-feedback (**Y**) from one neuron to cancel event-feedback (**Q**) from another, reducing the need for neuron-specific alignment. Independent noise disrupts these correlations, so cancellation requires the two pathways to align for the same downstream neuron. This suggests a functional role for intrinsic variability in promoting alignment of cortical feedback.

Next, we tested the effect of partitioning the apical compartment into multiple dendritic branches, in line with evidence that apical dendritic branches function as partially independent integration compartments ^32,33^. Each branch received event- and burst-feedback from distinct subsets of downstream neurons, such that burst-feedback plasticity had to cancel only the event-feedback within its corresponding branch. We found that increasing the number of branches accelerated **QY**-alignment (Fig. 3f). This suggests a functional role for dendritic branching in facilitating alignment of feedback pathways.

Finally, we examined whether **QY**-alignment remained robust in the presence of teaching signals (Fig. 3g). As expected, in the absence of teaching signals, **Y** plasticity drove near-perfect alignment with the fixed **Q** pathway (see Supplementary Information for analysis). Introducing teaching signals produced a modest increase in the final **QY** angle, because the apical deviations minimised by **Y** plasticity now reflected both pathway misalignment and teaching-driven feedback changes. Nevertheless, the final angles remained close to zero across teaching-signal strengths, indicating that local **Y** plasticity can maintain high feedback-pathway alignment even in the presence of task-related apical drive.

### Feedback-pathway alignment enables deep credit assignment

Having shown that local burst-feedback plasticity can bring the **Y** pathway into balance with the event-feedback pathway **Q**, we next asked whether this balance could be maintained during task learning. We therefore focused on a learning configuration with plastic feedforward weights **W** and burst-feedback weights **Y**, while keeping event-feedback weights **Q** fixed (Fig. 4a).

**Figure 4.**
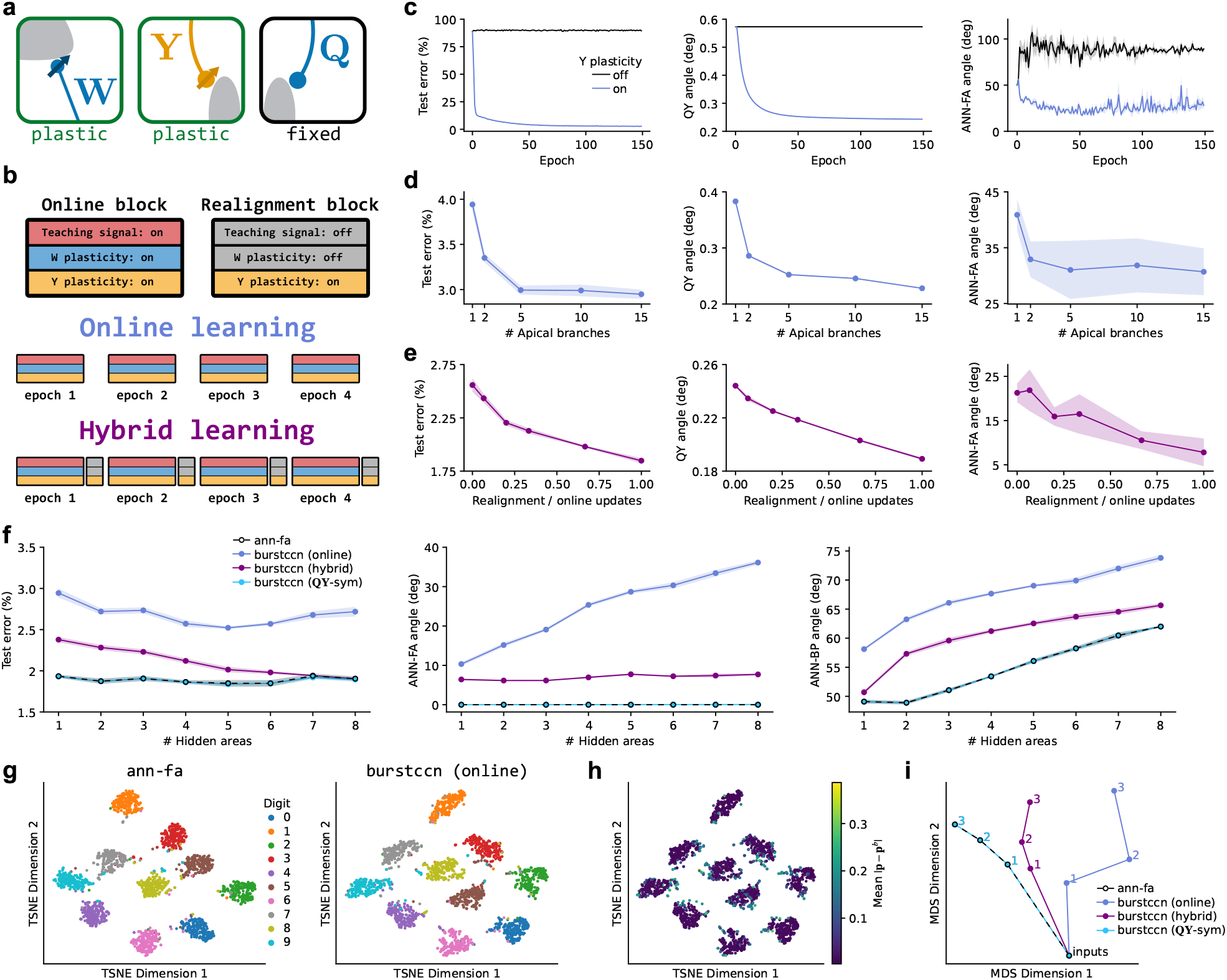
Feedback-pathway alignment enables deep credit assignment. **a**, Feedforward and burst-feedback plasticity configuration: feedforward weights **W** and burst-feedback weights **Y** are plastic, while event-feedback weights **Q** are fixed. **b**, Schematic of the two training block types and learning regimes. Online blocks involve learning with a teaching signal present and both **W** and **Y** plasticity enabled, whereas realignment blocks involve learning only the **Y** weights, without a teaching signal and with **W** plasticity disabled. In online learning, training consists of sequential epochs of online blocks, whereas hybrid learning includes a realignment block after each online block. **c**, Online BurstCCN training on MNIST with and without **Y** plasticity. Left: test error over training. Middle: alignment angle between **Q** and **Y** weights. Right: angle between the weight updates and the corresponding artificial neural network feedback-alignment (ANN-FA) updates. (**d**,**e**), Same metrics as in (**c**), shown for online BurstCCN with varying numbers of apical branches (**d**) and for hybrid BurstCCN with varying realignment block lengths (**e**). **f**, MNIST training comparison of ANN-FA and BurstCCN (online, hybrid, and **QY**-symmetric) models with varying numbers of hidden areas. Left: test error. Middle: angle between updates in the first hidden area and the corresponding ANN-FA updates. Right: angle between updates in the first hidden area and the corresponding backprop (ANN-BP) updates. **g**, t-SNE visualisations of neural responses to MNIST test images, coloured by class, in the second hidden area. Left: ANN-FA. Right: online BurstCCN. **h**, Same t-SNE visualisation as in (**g**, right), coloured by the mean change in burst probability across neurons relative to baseline. **i**, Representational trajectories for the models in (**f**), visualising the transformation of inputs through successive areas in a two-dimensional space using MDS. Numbers denote the corresponding hidden areas. Lines represent the mean, with standard error depicted by the shaded area (n = 5).

We directly compared BurstCCN feedforward weight updates to those computed for artificial neural networks (ANNs) using backpropagation (ANN-BP) or Feedback Alignment (ANN-FA) ^34^. ANN-BP computes exact task-error gradients using feedback weights precisely matched to those in the feedforward pathway, a requirement that is difficult to reconcile with biological circuits. ANN-FA avoids this requirement by using fixed random feedback weights and provides an analogue of the fixed **Q** feedback configuration used here.

To test whether **QY**-alignment could be maintained during task learning, we trained BurstCCN on a simple image recognition task (MNIST; see Methods) in two learning regimes: online and hybrid learning (Fig. 4b). Online learning consisted entirely of *online blocks*, in which task-related teaching signals were applied and both **W** and **Y** were plastic. In hybrid learning, each online block was followed by a *realignment block*, in which teaching signals were removed and plasticity was restricted to **Y**.

Rather than imposing perfect **QY**-symmetry, we introduced a small initial mismatch between the **Q** and **Y** pathways and compared online learning with an equivalent network in which **Y** was fixed (Fig. 4c). Fixing **Y** prevented the initial mismatch from being corrected, biasing apical feedback so that feedforward updates were not aligned with ANN-FA updates and task performance failed to improve. In contrast, with plastic **Y** weights, **QY**-alignment improved as the network learned the task, allowing it to achieve strong performance. We next varied the number of apical branches and found that additional branches improved both **QY**-alignment and task performance (Fig. 4d). With hybrid learning, longer realignment blocks improved **QY**-alignment and performance (Fig. 4e), consistent with stronger **QY**-alignment producing feedforward updates that were better aligned with ANN-FA updates (Fig. S3). Together, these results show that **Y** plasticity is critical for maintaining alignment during task learning.

We then examined how credit assignment scales with network depth. We compared ANN-FA with three BurstCCN configurations: an ideal setting with fixed **QY**-symmetry, the hybrid learning regime, and the online learning regime. Across all depths and learning regimes, BurstCCN consistently achieved strong performance below 3% test error (Fig. 4f). In networks with eight hidden areas, the ideal **QY**-symmetric configuration achieved 1.89 *±* 0.03% test error, comparable to both ANN-FA (1.90 *±* 0.03%) and hybrid BurstCCN (1.90 *±* 0.02%). Online BurstCCN also remained effective with this deep network, achieving 2.72 *±* 0.06% test error. Weight update alignment largely mirrored task performance, with better-performing models exhibiting closer alignment to ANN-FA and ANN-BP updates. These results show that BurstCCN can maintain effective credit assignment as network depth increases.

The close alignment between the **QY**-symmetric BurstCCN and ANN-FA updates is expected because these updates can be related analytically. In both ANN models and BurstCCN, feedforward weight updates involve a postsynaptic credit signal and presynaptic activity:

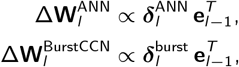

where **e**_*l*−1_ denotes presynaptic activity, corresponding to the event rate in BurstCCN. The term 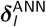 denotes either the exact error gradient derived from ANN-BP or the feedback error signal derived from ANN-FA. The term 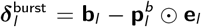 is the difference between the burst rate and its baseline. We show that perfect **QY**-symmetry enables BurstCCN feedforward updates to closely approximate ANN-FA updates and, importantly, ANN-BP updates when feedback weights are aligned with feedforward weights (see Supplementary Information for a detailed derivation).

Finally, we investigated whether the close update alignment resulted in similar neural representations across models. ANN-FA and online BurstCCN learned similar representations, forming clearly separated digit clusters when visualised using t-SNE (Fig. 4g). Within BurstCCN representations, input samples located further from their cluster centres exhibited larger changes in burst probability (Fig. 4h), predicting that ambiguous sensory inputs should evoke more heterogeneous bursting across cortical populations. To track representations across areas, we used Representational Trajectory Analysis (RTA) ^35^ to map successive area representations into a shared 2D space (Fig. 4i). As expected, the representational trajectories reflected the alignment between ANN-FA and the different BurstCCN variants (cf. Fig. 4f).

Together, these results show that local apical plasticity can maintain **QY**-alignment during task learning, enabling BurstCCN to perform effective credit assignment across deep hierarchical cortical networks.

### BurstCCN scales to complex image recognition and reinforcement learning tasks

Having demonstrated that BurstCCN can perform effective credit assignment on MNIST, we next asked whether it scales to more challenging tasks. We therefore evaluated BurstCCN on complex image-recognition and reward-based navigation tasks (Fig. 5). To support learning in these more challenging tasks, we introduced a plastic-feedback configuration in which both feedback weights, **Y** and **Q**, were updated (Fig. 5a; see Methods). This allowed these feedback weights to align more closely with the feedforward weights, enabling BurstCCN to better approximate ANN-BP updates.

**Figure 5.**
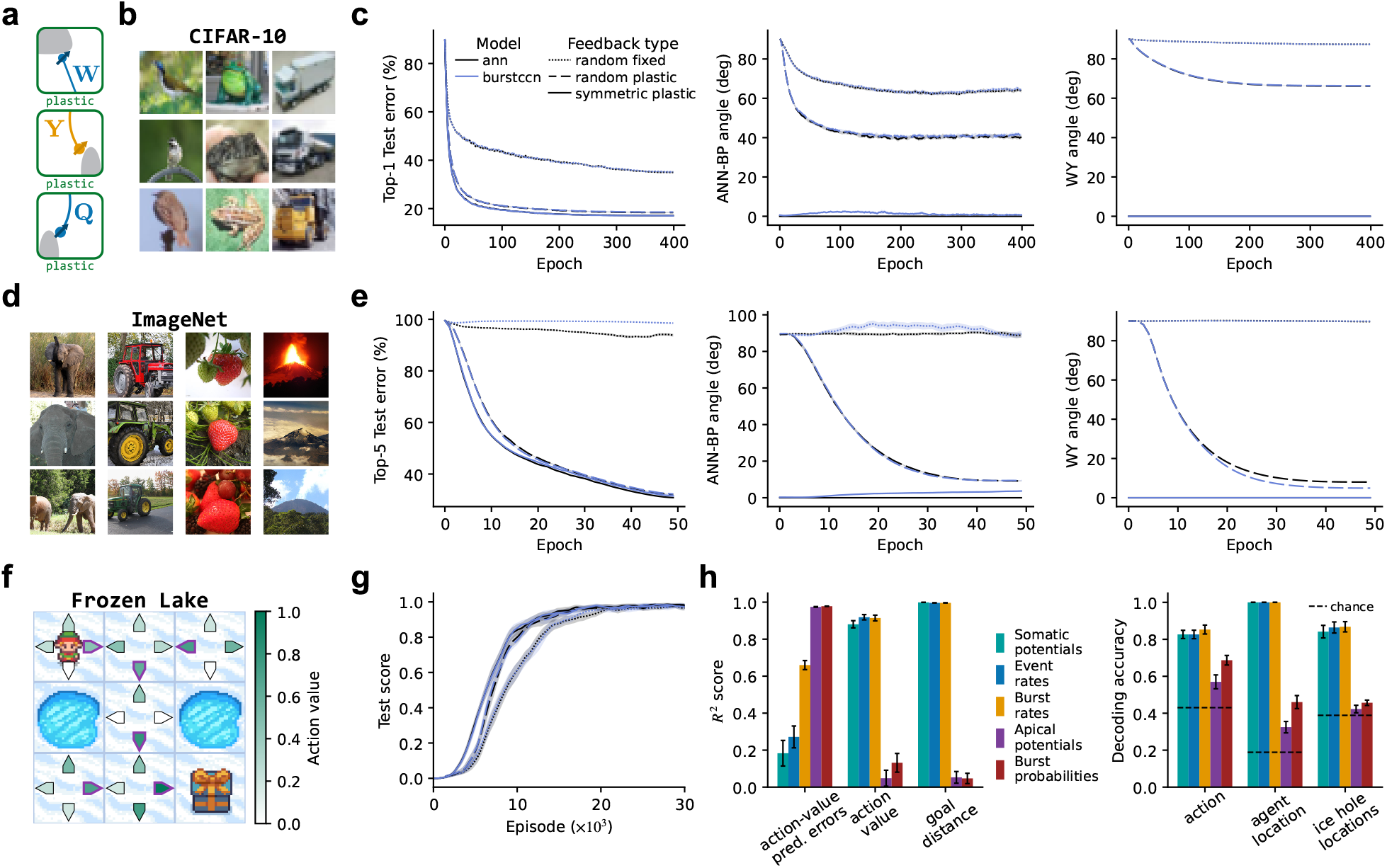
BurstCCN learns complex image recognition and reinforcement learning tasks. **a**, Fully plastic configuration: feed-forward weights **W** and both feedback pathways, **Q** and **Y**, are plastic. **b**, Example images from the CIFAR-10 dataset. **c**, CIFAR-10 training results comparing convolutional ANN and **QY**-symmetric BurstCCN models with different feedback (both **Q** and **Y**) configurations: random fixed feedback weights, random plastic feedback weights, and symmetric plastic feedback weights. Left: Top-1 test error over training. Middle: Angle between weight updates and backprop updates. Right: Angle between feedforward (**W**) and feedback (**Y**) weights. **d**, Example images from the ImageNet dataset. **e**, ImageNet training results, shown as in (**c**). Left: Top-5 test error. Middle: Alignment angle with backpropagation updates. Right: Angle between **W** and **Y** weights. **f**, Schematic of the Frozen Lake reinforcement learning task. The agent learns to navigate to the reward while avoiding holes in the ice. Purple outlines indicate the maximum learned action value (Q-value) in each state, corresponding to the optimal policy in this example. **g**, Task performance on Frozen Lake, measured as the proportion of successful episodes. **h**, Decoding of task-relevant variables from internal network quantities. Left: decoder *R*^2^ score for continuous variables, including action-value prediction error, the learned value of the selected action and the distance to the goal. Right: decoder accuracy for categorical variables, including the selected action, the agent’s location and the locations of ice holes within the map. Lines represent the mean across runs, with shaded areas indicating standard error (n = 5). The images in (b) and (d) were obtained from publicly available datasets (CIFAR-10 and ImageNet).

We first evaluated BurstCCN on the CIFAR-10 image classification benchmark (Fig. 5b), using a convolutional architecture (Fig. 5c; see Methods). We compared fixed random feedback with two plastic-feedback variants: random-plastic feedback, which was also initialised randomly, and symmetric-plastic feedback, which was initialised symmetrically with the feedforward pathway. As expected, fixed random feedback was sufficient for learning but showed the largest misalignment with ANN-BP updates and the worst performance. In contrast, random-plastic feedback substantially improved performance from the same random initialisation, approaching the symmetric-plastic variant in both test error and ANN-BP update alignment. The symmetric-plastic BurstCCN produced the lowest test error (17.03 *±* 0.21%) and closest alignment with ANN-BP updates, performing comparably to the corresponding symmetric-plastic ANN (17.10 *±* 0.11%). Importantly, across all three configurations, BurstCCN achieved performance comparable to an ANN with the same architecture.

We next evaluated BurstCCN on ImageNet, a larger and more challenging visual-recognition benchmark (Fig. 5d; see Methods). Fixed random feedback failed to learn (Fig. 5e), consistent with the poor scaling of random feedback in deep convolutional networks ^36^. By contrast, feedback plasticity enabled BurstCCN to achieve performance close to the symmetric-plastic configuration, despite starting with a random feedback initialisation. In the symmetric-plastic configuration, BurstCCN achieved 31.34 *±* 0.05% top-5 test error, close to the ANN with the same architecture (30.66 *±* 0.03%). This improvement coincided with increased alignment between feedforward and feedback pathways, indicating that adaptive cortical feedback plasticity is important for scaling to more demanding recognition tasks.

Finally, we tested whether BurstCCN can learn without explicit supervised targets. We trained the network in a DQN-style reinforcement learning setup ^37^ on the Frozen Lake navigation task, where the agent must reach a goal location while avoiding ice holes and receives only sparse rewards upon success (Fig. 5f; see Methods). Using the same feedback configurations as in the image-classification experiments, BurstCCN learned the task even with fixed random feedback, but learned fastest with the symmetrically initialised configuration (Fig. 5g).

To examine what task-relevant information is represented by the cortical network during reinforcement learning, we trained linear decoders to predict task- and value-related variables from internal network signals (Fig. 5h). Feed-forward quantities, including somatic potentials and event rates, supported decoding of the selected action, its action value and task-state information, including the agent’s location, ice-hole locations and distance to the goal. By contrast, feedback-related quantities, including apical potentials and burst probabilities, primarily supported decoding of action-value prediction errors. Burst rates, which combine event-rate and burst-probability information, predicted both task-state variables and action-value prediction errors. Together, these results predict that task state and action values are represented in feedforward somatic and event-related signals, whereas prediction-error signals are encoded in feedback-related apical potentials and burst probabilities.

### Cell-type-specific pathways account for plasticity, circuit architecture and learning observations

Thus far, synapses in BurstCCN have been allowed to take positive or negative values, such that individual connections could provide either excitation or inhibition. However, biological neurons obey Dale’s law, constraining each neuron to exert either excitatory or inhibitory effects on its postsynaptic targets. We therefore constructed a Dalean BurstCCN in which pyramidal neurons provide only excitatory outputs and inhibition is mediated by four interneuron types expressing parvalbumin (PV), somatostatin (SST), vasoactive intestinal peptide (VIP) or neuron-derived neurotrophic factor (NDNF) (Fig. 6a; see Methods). The feedforward, event-feedback and burst-feedback pathways were decomposed into excitatory and inhibitory components consistent with cortical microcircuits: **W**^+^ and **W**^−^ correspond to pyramidal excitation and PV-like somatic inhibition, **Q**^+^ and **Q**^−^ to excitatory feedback projections and NDNF-like apical inhibition, and **Y**^−^ and **Y**^+^ to SST-like apical inhibition and VIP-mediated disinhibition. This decomposition preserves the functional roles of the original **W, Q**, and **Y** pathways and is consistent with known interneuron connectivity and pathway-specific short-term synaptic dynamics ^43–46^. We next show that the predictions arising from this variant of the model account for several experimental observations.

**Figure 6.**
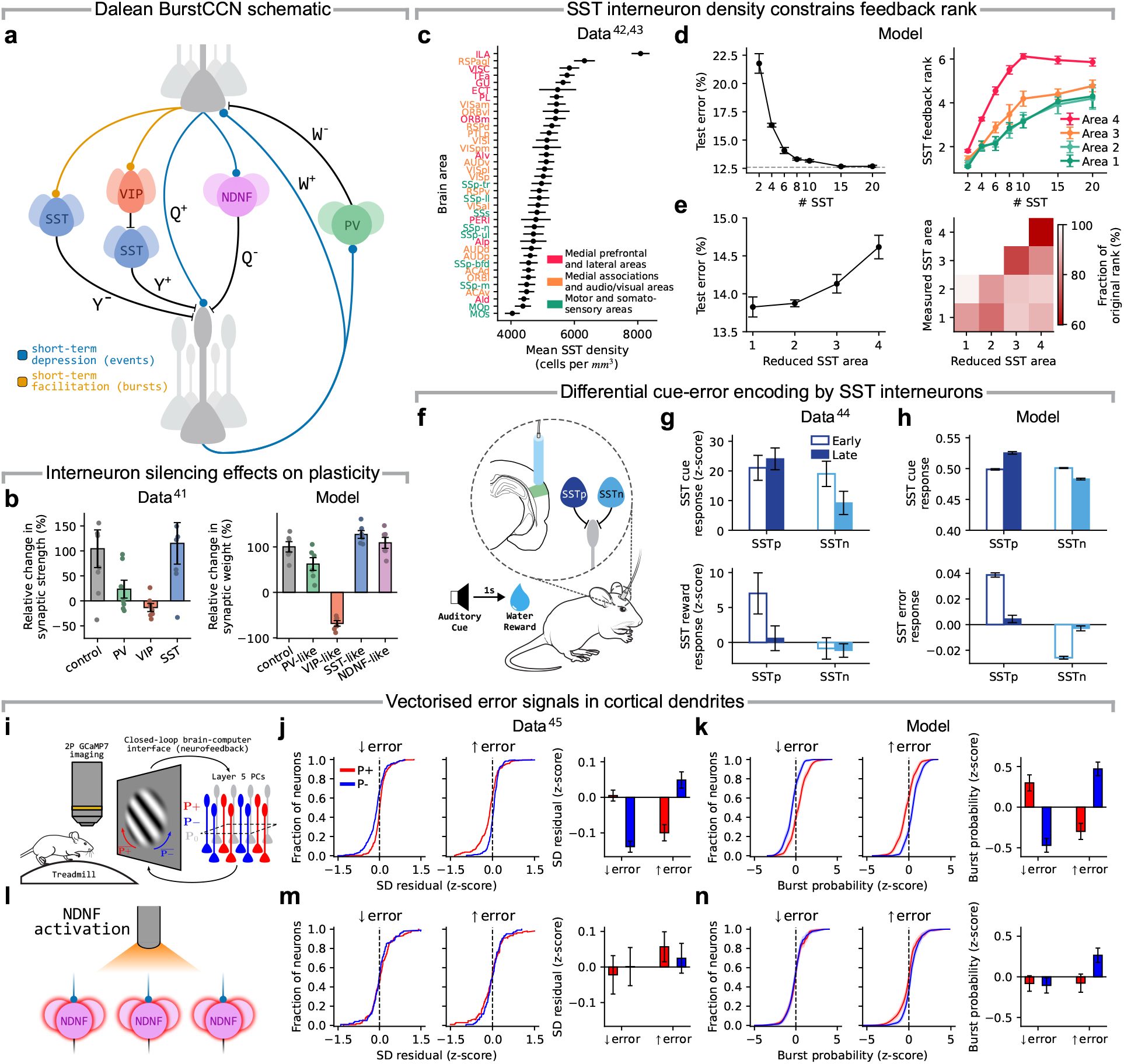
Cell-type-specific pathways account for plasticity, circuit architecture and learning observations. **a**, Model schematic of the Dalean BurstCCN, incorporating multiple interneuron types expressing parvalbumin (PV), somatostatin (SST), vasoactive intestinal peptide (VIP), or neuron-derived neurotrophic factor (NDNF). Each pathway in the standard BurstCCN is decomposed into excitatory (or disinhibitory, in the case of VIP–SST feedback) and inhibitory components. **b**, Left: Experimental data showing how suppression of different interneuron types shapes synaptic plasticity in local circuits ^38^. Cortical synaptic efficacy was measured using layer 4-evoked postsynaptic potential amplitudes in layer 2/3 pyramidal neurons following electrical stimulation of layer-4 input together with optogenetic stimulation of feedback. Right: Model results from the Dalean BurstCCN, showing relative changes in feedforward synaptic weights, quantified as Δ**W**_1_*/***W**^init^, when individual interneuron populations are selectively silenced. **c**, Mean SST interneuron density across mouse cortical areas, ordered from highest to lowest mean SST density. Adapted from Kim et al. ^39^, Wang and Yang ^40^ . **d**, Left: F-MNIST test error of the Dalean BurstCCN as the number of SST interneurons per area was varied equally across all areas. Dashed line indicates test error with full-rank apical feedback in each area. Right: Apical dendritic feedback rank in each area for the same networks. **e**, Left: F-MNIST test error for networks trained with a reduced number of SST interneurons in a single area, with all other areas unchanged. Right: Apical feedback rank in each area for the same single-area reductions, expressed as a fraction of the corresponding rank for that area in a network without any reduction in SST interneurons. **f**, Schematic of fibre photometry recordings from Chevy et al. ^41^, distinguishing reward-responsive (SSTp) and reward-non-responsive (SSTn) SST interneurons in an associative auditory Pavlovian task. **g**, Quantification of experimentally observed SSTp and SSTn fluorescence changes during early and late training. Top: cue response, quantified as the change in fluorescence from cue onset to cue offset. Bottom: reward response, quantified as the change in fluorescence from reward onset to 1s after reward onset. **h**, Quantification of SSTp and SSTn activity in the model during stimulus and error presentation. Top: mean activity during image presentation. Bottom: error response, computed as the change in activity between stimulus presentation and target (error) presentation. **i**, Schematic of the closed-loop BCI task from Francioni et al. 42 , in which two-photon imaging of layer 5 pyramidal neurons in mouse retrosplenial cortex is used to control the rotation of a Gabor grating. The grating angle was determined by the difference in average activity between P+ and P− populations (P+ − P−): increased P+ activity drove clockwise rotation towards 90°, whereas increased P− activity drove anticlockwise rotation towards 0°. **j**, Experimental data reproduced from Francioni et al. 42 , showing cumulative distributions of somato-dendritic residuals for P+ and P− neuron populations during periods of decreasing and increasing task error, z-scored across all neurons. Left: decreasing error. Middle: increasing error. Right: Mean z-scored somato-dendritic residuals for P+ and P− neurons during both error conditions. **k**, Model (Dalean BurstCCN): cumulative distributions of burst probability for P+ and P− neuron populations in response to positive and negative feedback teaching signals, corresponding to decreasing and increasing task errors, respectively. Burst probabilities are represented as z-scores across all neurons. Left: positive teaching signal. Middle: negative teaching signal. Right: Mean z-scored burst probability for P+ and P− neurons during both teaching signal conditions. **l**, Experimental schematic from Francioni et al. ^42^ showing optogenetic activation of layer 1 NDNF interneurons during the BCI task. **m**, Experimental data reproduced from Francioni et al. ^42^ , showing the same comparison as in **j** during optogenetic activation of layer 1 NDNF interneurons. **n**, Model (Dalean BurstCCN), showing the same burst-probability comparison as in **k**, but with a constant positive input to NDNF interneurons.

#### Cell-type-specific control of feedforward synaptic plasticity

Experiments show that suppressing distinct interneuron populations differentially affects pyramidal-cell synaptic plasticity ^38^ (Fig. 6b, left). We selectively silenced each interneuron type in the model while the network learned to increase pyramidal-cell activity towards a target (see Methods). Silencing SST or NDNF interneurons increased dendritic bursting and strengthened potentiation, whereas silencing VIP interneurons increased SST activity, strengthening inhibition of apical dendrites and shifting plasticity towards depression. Finally, silencing PV interneurons increased somatic activity, reducing the additional feedforward potentiation required to reach the target (Fig. 6b, right; Fig. S4). Together, these model perturbations match the direction of the experimentally observed effects of interneuron-specific manipulations on pyramidal-cell plasticity ^38^.

#### SST density constrains the cortical feedback rank

Anatomical studies show that SST interneuron density increases along the cortical hierarchy ^39,40^ (Fig. 6c). Since SST interneurons mediate burst-related teaching signals in the Dalean BurstCCN, we expected the size of this population to constrain the dimensionality of feedback error signals. To test this, we trained the model on the F-MNIST image recognition task with different SST population sizes across areas (see Methods). Reducing SST population size uniformly across areas imposed a bottleneck on the burst-mediated teaching signal, reducing its rank and impairing learning (Fig. 6d). When SST population size was reduced in individual areas, reductions in later areas produced the strongest performance impairment (Fig. 6e), indicating that preserving the dimensionality of burst-mediated feedback becomes increasingly important deeper in the hierarchy, where flexible representations place greater demands on learning signals ^47,48^. Consistent with this, alignment with full-rank ANN updates showed a similar dependence on SST population size, decreasing with smaller populations (Fig. S5). These results provide a functional interpretation of the observed cortical SST gradient, providing the key prediction that SST interneurons limit the dimensionality of error-related signals in the cortex.

#### Differential encoding of cue and error signals in SST subpopulations

Recent work identified two SST subpopulations during reward learning: reward-responsive SST neurons (SSTp) and non-responsive SST neurons (SSTn) ^41^ (Fig. 6f). Across learning, SSTp neurons increased cue responses and suppressed reward responses, whereas SSTn neurons decreased cue responses and remained weakly responsive to reward (Fig. 6g). We expected analogous subpopulation-specific changes to emerge in the model by treating stimulus-related SST activity as a proxy for cue responses and error-related SST activity as a proxy for reward-related responses. In line with the experimental methods ^41^, we divided model SST neurons into SSTp and SSTn subpopulations based on their pre-learning error responses (see Methods). SSTp neurons increased stimulus-driven activity and reduced their error response across learning, whereas SSTn neurons decreased stimulus-driven activity and showed weaker negative error responses (Fig. 6h). These learning-dependent changes naturally arise in the model SST interneurons because they decode burst-related feedback signals that predict the direction of subsequent synaptic plasticity.

#### Vectorised dendritic instructive signals during BCI learning

Recent closed-loop brain–computer interface (BCI) work shows that cortical pyramidal neurons receive neuron-specific error signals during learning ^42^. In this task, mice controlled the rotation of a visual stimulus through two predefined layer 5 pyramidal-cell populations, P+ and P−, which had opposing causal roles. Increasing P+ activity drove clockwise rotation, whereas increasing P− activity drove anticlockwise rotation (Fig. 6i). Somato-dendritic residuals, defined as dendritic activity unexplained by somatic activity, differed systematically between these populations depending on whether task error was decreasing or increasing (Fig. 6j). We implemented an analogous BCI setup, partitioning pyramidal neurons in the model into P+ and P− populations and quantifying neuron-specific error signals as deviations in burst probability from baseline. The model reproduced the experimental pattern: teaching signals corresponding to decreasing error elevated burst probability in P+ relative to P− neurons, whereas teaching signals corresponding to increasing error reversed this relationship (Fig. 6k). As expected, this shows that BurstCCN generates neuron-specific feedback signals aligned with each neuron’s causal role in the task. The model further predicts that disrupting apical feedback pathways should impair this relationship. Consistent with this, optogenetic activation of NDNF interneurons in the BCI task decoupled somato-dendritic residuals from the error direction ^42^ (Fig. 6l,m). Similarly, NDNF activation in our model eliminated the relationship between burst probability and error direction (Fig. 6n). Together, these results support burst probability as a candidate dendritic signal for credit assignment during in vivo learning.

## Discussion

Through AI-inspired computational modelling, we have shown that differential cortical feedback can drive dendrite-dependent, burst-mediated synaptic plasticity across multiple levels of cortical processing, providing a mechanistic solution to hierarchical credit assignment ^1–3^. Previous approaches to biologically plausible credit assignment have shown how multi-compartment neurons can encode learning signals, but remain limited in their ability to propagate error signals through deep cortical hierarchies and often require multi-phase learning ^5,11^. By using pathway-specific feedback to multiplex event- and burst-related signals, our model addresses these limitations and enables learning in complex recognition and reward-driven tasks. The framework predicts that learning should depend on cell-type-specific feedback to regulate dendritic excitation–inhibition balance, shape neuron-specific error signals, and coordinate plasticity across cortical hierarchies. Together, these results suggest that distinct cortical cell types jointly coordinate hierarchical learning in the cortex.

Distal dendrites are well positioned to integrate cortical feedback and transform it into local instructive signals for plasticity. A fundamental prediction of our model is that apical dendrites encode neuron-specific error signals that are distinct from the information represented at the soma, in line with recent observations ^33,42,49^. Our model further predicts that dendritic error encoding depends on two cortical feedback pathways with distinct short-term plasticity dynamics, which provide separate event- and burst-related feedback signals. The simultaneous availability of these signals at distal dendrites enables learning without distinct temporal phases. Effective dendritic error encoding requires these two feedback pathways to be aligned, such that dendritic excitation–inhibition balance is maintained in the absence of error, while deviations from this balance generate error signals. Furthermore, we show that intrinsic neural variability, dendritic branching, and periods without teaching signals promote this alignment, suggesting functional roles for these biological features and circuit states in supporting subsequent learning.

Beyond pyramidal-cell computations, our results highlight how interneuron diversity can coordinate brain-wide credit assignment. By targeting apical dendrites, SST interneurons provide the inhibitory control required to maintain dendritic excitation–inhibition balance and allow burst-dependent error signals to be selectively isolated during learning. This is consistent with known error-related functions of SST interneurons ^50^, evidence that SST interneurons gate top-down inputs during associative learning ^51^, and findings that thalamocortical bursts preferentially recruit dendrite-targeting SST cells over PV cells to regulate dendritic integration ^52^. Anatomically, SST interneuron density increases along the cortical hierarchy ^39,40^. Our model assigns this gradient a functional role by predicting that higher SST density supports higher-dimensional feedback in higher-order association areas, where flexible representations place greater demands on feedback dimensionality. Together, these findings suggest that specialised SST connectivity goes beyond circuit stabilisation ^27^, serving as a core component of deep credit assignment.

Our framework also raises several questions. Although we show that BurstCCN supports learning driven by internally generated targets (Fig. 2), externally provided labels (Figs. 4 and 5), and sparse rewards (Fig. 5), it remains unclear where the corresponding teaching signals are computed in the brain. Our model predicts that such signals should modulate apical dendrites, motivating future work to determine whether subcortical structures such as thalamic nuclei or the dopaminergic system perform such control ^53,54^. In addition, although short-term plasticity (STP) has been shown to decode events and bursts ^18,22^, here we assumed idealised STP-based decoding. Future work should test the robustness of this mechanism under noisy and heterogeneous STP dynamics. Furthermore, our spiking model implementation includes a number of biophysical properties, but real pyramidal neurons possess richer active dendritic properties, including localised NMDA spikes and complex calcium dynamics. Determining whether this biological complexity enhances signal separation or introduces disruptive noise is important for developing a biophysical account of credit assignment. More broadly, extending the model to incorporate temporal credit assignment ^2,55^ is essential for understanding how the cortex assigns credit across time.

In conclusion, our results show that cell-type-specific control of apical dendrites provides a biologically grounded solution to the hierarchical credit assignment problem. This framework recasts cortical cell diversity as a fundamental requirement for learning complex tasks rather than mere anatomical complexity.

## Acknowledgements

We would like to thank the Neural & Machine Learning group, Jack Mellor, Quentin Chevy and Ian Cone for useful feed-back. WG was funded by an EPSRC Doctoral Training Partnership award (EP/R513179/1), HWZ by the Wellcome Trust (Neural Dynamics PhD Program) and RPC by the Medical Research Council (MR/X006107/1), BBSRC (BB/X013340/1) and an ERC-UKRI Frontier Research Guarantee Grant (EP/Y027841/1). This work used the HPC system BluePebble at the University of Bristol, UK.

## Author contributions

The author contributions are as follows:

Conceptualisation: WG, HWZ and RPC. Methodology: WG, HWZ, and RPC. Simulations: WG, HWZ (initial results for MNIST, CIFAR-10, dendritic plasticity, RL, Spiking BurstCCN, Continuous BurstCCN, Dalean BurstCCN), AD (initial interneuron reduction analysis), PM (initial cell-type plasticity simulations) and KN (initial deep RL simulation). Analytical results: WG and JP. Investigation: WG, HWZ, and RPC. Visualisation: WG, HWZ, and RPC. Supervision: RPC. Writing: WG, HWZ, and RPC.

## Methods

### Bursting Cortico-Cortical Networks

Across experiments, we used multiple implementations of BurstCCN, differing in their level of abstraction and biological detail. This allowed us to relate the model to biophysical mechanisms while using more abstract implementations for more complex tasks. We first describe the burst-dependent plasticity rule used in the spiking model. We then focus on the discrete-time, rate-based implementation of BurstCCN, which provides a clearer description of the key model components and their relationship to credit assignment (see Supplementary Methods for the spiking, continuous-rate and Dalean variants of the model).

#### Spike-based burst-dependent plasticity rule

Here we describe the learning rule used by the spiking BurstCCN (see Supplementary Methods for details on this model). The feedforward **W** weights in the spiking BurstCCN remained plastic throughout stimulus presentation. Building on the burst-dependent plasticity rule introduced in Burstprop ^11^, these weights were updated according to:

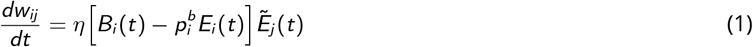

where *η* controls the learning rate, 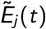 is a moving average of the presynaptic neuron’s event rate, and 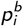 is a fixed baseline burst probability. Postsynaptic burst events increase the synaptic weight in proportion to 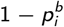, whereas isolated single-spike events decrease it in proportion to 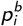. As a result, neurons with burst probability greater than 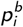 undergo LTP, whereas those with lower burst probability undergo LTD.

#### Discrete-time rate-based BurstCCN

The discrete-time, rate-based implementation of BurstCCN abstracts the spiking and continuous-time rate-based models (see Supplementary Methods). Rather than modelling individual spiking neurons, it abstracts the population-level dynamics of ensembles of these pyramidal neurons. It further abstracts the continuous temporal dynamics by using discrete timesteps, in which input–output example pairs are presented to the network and processed through a feedforward pass followed by a feedback pass.

For each example, the feedforward pass begins by encoding the input through the event rates of input area ensembles, **e**_0_. These event rates propagate through the short-term depressing (STD) feedforward synaptic weights, **W**_*l*_, at each area *l* = 1, …, *L*, producing the somatic potentials of the next area, **v**_*l*_ = **W**_*l*_ **e**_*l*−1_. Feedforward bias terms are also included in the calculation of somatic potentials but are omitted from the notation for simplicity. The ensemble-level event rates are then computed by applying a nonlinear activation function, *f*, to the somatic potentials, **e**_*l*_ = *f* (**v**_*l*_ ). This process is repeated for all subsequent areas until the output area is reached, equivalent to the function computed by a standard feedforward artificial neural network.

In the feedback pass, the output area event rates, **e**_*L*_, are compared with the desired output targets, **e**_target_, to produce a signed error used to define a teaching signal, **i**_teacher_ = *κ*(**e**_target_ − **e**_*L*_), where *κ* is the teaching signal strength. This teaching signal provides error information that is propagated backward through the network, altering apical compartment potentials and, as a result, ensemble-level burst probabilities. The burst probabilities are first computed at the output area directly as:

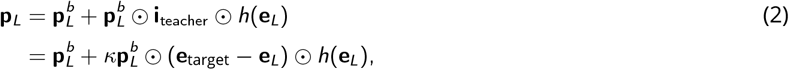

where ⊙ denotes the element-wise product. In this equation, 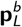 represents a vector of baseline burst probabilities for the output area ensembles, reflecting their burst probability in the absence of a teaching signal. To simplify the notation here, and for all experiments, we assume a homogeneous baseline within each area, such that 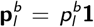 where 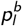 is a scalar baseline burst probability for area *l* and **1** is a vector of ones with the same dimension as **p**_*l*_ . The function, 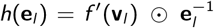, incorporates the activation function derivative into the feedback pathway, ensuring a correspondence to the error gradients computed by the backprop algorithm (see Supplementary Information for the link to backprop). Its inclusion of 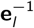 serves to cancel with a subsequent multiplication by **e**_*l*_ in the calculation of the ensemble-level burst rates, **b**_*l*_ = **e**_*l*_ ⊙ **p**_*l*_, derived from the definition of burst probability, 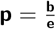.

Following the calculation of the burst rates in a given area *l* + 1, they are then decoded and sent back to the apical dendrites of area *l* by a set of STF feedback weights, **Y**_*l*_ . Additionally, a novel set of apical dendrite-targeting STD feedback weights, **Q**_*l*_, are used to decode and send back event rates. Similar to the feedforward connections, both feedback types are assumed to decode their respective signals exactly and without interference. Together, the two types of feedback determine the apical potentials of area *l* :

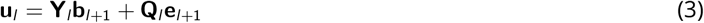

which can be understood as a weighted (through **Y**_*l*_ and **Q**_*l*_ ) combination of error-dependent burst signals, **b**_*l*+1_, and error-invariant event signals, **e**_*l*+1_. The ideal configuration for the weights of these connections is such that these two signals precisely cancel each other when the ensembles in the next area are at baseline bursting activity 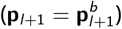, ensuring that the apical potentials in the current area are silenced (**u**_*l*_ = 0). Specifically, this cancellation is achieved when the weights satisfy the condition, 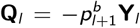, referred to as the **QY**-symmetric state. Intuitively, since the burst probability is between 0 and 1, the **Q**-weights will be smaller than their corresponding **Y**-weights in this state, compensating for the higher frequency of events relative to bursts. For all areas prior to the output area, the apical potentials directly control the ensemble-level burst probabilities as follows:

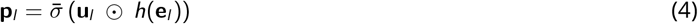

where 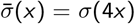 is a scaled sigmoid function. This scaling gives the feedback nonlinearity an approximately unit slope around zero, avoiding implicit downscaling of feedback signals. The same process is repeated for every area, in reverse order, to compute their apical potentials and ensemble burst probabilities.

The feedback-induced changes to burst probabilities across all areas represent a credit assignment process with respect to a mean-squared error (MSE) defined at the output area, 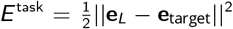. These credit signals are used to update the feedforward weights using a burst-dependent synaptic plasticity rule:

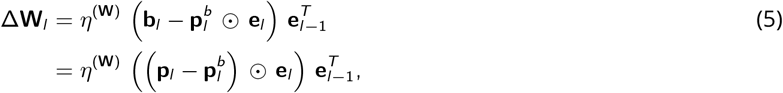

where *η*^(**W**)^ is a learning rate and ·^*T*^ indicates the transpose operation. The term 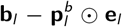 corresponds to the postsynaptic change in burst rate from its baseline value. Equivalently, this can be written as 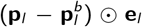, the deviation in burst probability from baseline weighted by the postsynaptic event rate. This represents a postsynaptic credit signal that is assigned to individual feedforward synapses by multiplication with the presynaptic event rates, 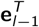. Feedforward biases were updated by applying the same learning rule with the presynaptic event rate set to 1.

In the absence of teaching signals, where **i**_teacher_ = 0, it is important that all pyramidal ensembles in all areas burst at their baseline probabilities to avoid unwanted changes to the feedforward **W**-weights. At the output area, this condition is satisfied by the definition of the burst probabilities in Equation 2. However, in the hidden areas, the signals to the apical compartments must cancel, which requires all feedback weights to be in the **QY**-symmetric configuration. To address the biological implausibility of the **Y** synapses having direct knowledge of **Q**, we introduce a learning rule for **Y**, inspired by earlier work ^5,28^, to learn this cancellation:

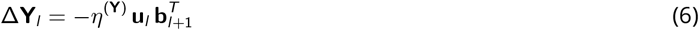

where *η*^(**Y**)^ is a learning rate. This learning rule explicitly aims to silence the apical potentials by aligning **Y** closer to the **QY**-symmetric configuration. In fact, in the Supplementary Information we prove that, under some reasonable assumptions, and in the absence of teaching signals at the output area, all **Y** weights will eventually converge to their optimal values, achieving **QY**-symmetry.

The following extensions were applied to the discrete-time rate-based model in specific experiments:

#### Somatic noise

To introduce independent variability in neuronal activity, we added Gaussian noise directly to the somatic compartment of each area. Specifically, for area *l*, the somatic potentials were computed as

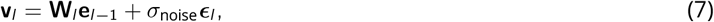

where ***ϵ***_*l*_ ∼ *N* (**0, I**) is a vector of independent Gaussian random noise and *σ*_noise_ denotes the standard deviation of the noise. Noise was sampled independently and identically distributed across neurons and across samples at each timestep.

#### Multiple apical dendritic branches

We extended the discrete-time BurstCCN by partitioning the apical compartment of each pyramidal cell into multiple branches to examine how this affects the ability to learn feedback-pathway alignment. For each pyramidal cell in area *l*, feedback inputs from area *l* + 1 were partitioned across *M*_*l*_ apical branches, with **Q** and **Y** connections from the same presynaptic source assigned to the same branch. For branch *m*, 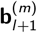 and 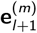 are the burst-rate and event-rate inputs received from area *l* + 1, and 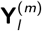 and 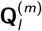 are the corresponding branch-specific feedback weights. The apical potential of branch *m* is given by

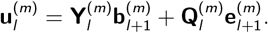

This allows plasticity of **Y** to be applied independently within each branch according to

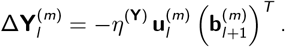

The total apical potential is then given by summing across branches:

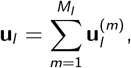

which is equivalent to the apical potential in the single-branch model (Equation 3).

#### Kolen–Pollack feedback learning

For the complex image classification experiments, fixed random feedback weights are known to be insufficient to support effective learning ^36^. We therefore allowed both feedback pathways, **Y** and **Q**, to learn using a Kolen–Pollack update rule ^56^, which provides a mechanism to align with the feedforward weights, **W**. The basic idea is that feedback plasticity in both pathways is driven by the same event- and burst-dependent signals as the forward pathway, causing them to undergo equivalent synaptic weight updates. With weight decay on all synaptic weights, the mismatches between differently initialised weights decrease over time, increasing alignment without the need for weight transport.

The burst-feedback weights were updated according to

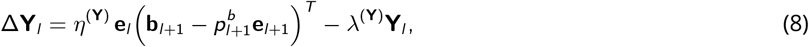

while the event-feedback weights were updated according to

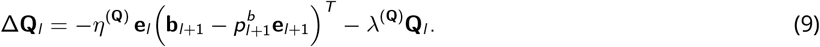

Here, *η*^(**Y**)^ and *η*^(**Q**)^ are feedback learning rates, and *λ*^(**Y**)^ and *λ*^(**Q**)^ are weight decay coefficients. For convolutional areas, the outer products above are replaced by the corresponding convolutional kernel-update operators. Importantly, we set 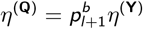 and *λ*^(**Q**)^ = *λ*^(**Y**)^ such that, when the feedback pathways are in the **QY**-symmetric configuration, this symmetry is preserved after each synaptic weight update across learning.

#### Alignment metrics

To quantify alignment between two sets of matrices, 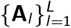 and 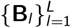, we first concatenated their vectorised forms across areas,

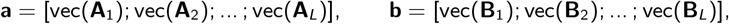

and then defined the alignment angle as

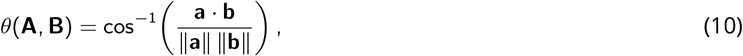

Using this definition, **QY**-alignment was computed from **A**_*l*_ = **Q**_*l*_ and **B**_*l*_ = −**Y**_*l*_, and **WY**-alignment from **A**_*l*_ = **W**_*l*_ and **B**_*l*_ = **Y**_*l*_ . Alignment to backpropagation (ANN-BP) and Feedback Alignment (ANN-FA) was computed in the same way, using the corresponding feedforward weight-update matrices in place of **A**_*l*_ and the reference ANN-BP or ANN-FA update matrices in place of **B**_*l*_ .

### Task details

#### Stationary target prediction task

The model hyperparameters used for this task were selected to best illustrate the dynamics of each model component and are provided in Table S2. The network consisted of two cell ensembles, each consisting of 100 neurons, connected in series as shown in Figure 1c. A fixed sensory input is provided to the somatic compartments of the first population through the synaptic weights, **W**_(1)_, which are free to learn following the burst-dependent plasticity rule in Equation 1. The second population receives the feedforward events of the first population into their somatic compartments through the synaptic weights, **W**_(2)_, which remain fixed throughout the entire task. The event rates of this second population are considered to be the output of the network.

Each stage begins with a 10s period in which no teaching signal is provided so that the network can reach an equilibrium before learning. This is followed by a 20s period in which a teaching signal is given to the second population’s apical compartments based on the difference between each neuron’s event rate and the target event rate. Finally, each stage ends with another 10s period without a teaching signal to test whether the network retains its most recently learned target.

#### Continuous nonlinear regression tasks

Similar to prior work ^5^, we constructed each nonlinear regression task using a randomly initialised fixed target network with the same architecture as the predictor network learning the task (Fig. 2a). A time-varying input, *x* (*t*), was given to both networks, and the target output, *y* (*t*), was defined by the event rates in the output area of the target network. In the brain, this task can be interpreted as a cortical area learning to transform a shared input into the activity pattern of another cortical area. For both tasks, the time constants used in the simulation for the somatic potentials, dendritic potentials and synaptic weights were set to *τ*_*v*_ = 0.1s, *τ*_*u*_ = 0.1s and *τ*_*W*_ = 100.0s, respectively.

##### Sinusoidal inputs

This task consisted of three sinusoidal inputs, *x*_*i*_ (*t*) = cos(2*πα*_*i*_ *t* + *β*_*i*_ ), with frequencies *α*_*i*_ ∼ *U*(0, 2.5) Hz and phase offsets *β*_*i*_ ∼ *U*(0, 2*π*) (Fig. 2b). For this task, both the predictor and fixed target networks used two-area architectures with 25 hidden units and a single output unit. The weights in the target network were initialised using Xavier normal initialisation with a gain of 3.

##### CatCam inputs

The CatCam dataset ^24^ is a collection of greyscale videos captured by a head-mounted camera on a cat exploring outdoor environments. Each video consisted of a finite number of frames, which were converted to a continuous-time input using the frame interpolation approach described by Reda et al. ^57^, enabling the video to be sampled at any temporal resolution. Each 320 *×* 240 pixel frame was cropped to a random 32 *×* 32 window and flattened into an input vector of length 1024. Videos were shown for their full duration, after which the next video immediately began with a new randomly selected crop location. After the final video, the input looped back to the start of the first video. For this task, both the predictor and fixed target networks used three-area architectures with 500 hidden units in each hidden area and 10 output units. A larger number of output units was used to provide a higher-dimensional teaching signal to the predictor network. The weights in the target network were initialised using Xavier normal initialisation with a gain of 2.

#### Feedback-only learning experiments

In these experiments, we tested whether local plasticity in the burst-feedback pathway could align the burst-feedback weights, **Y**, with fixed event-feedback weights, **Q**, while feedforward weights, **W**, remained unchanged. All experiments used the discrete-time rate-based BurstCCN. Only the **Y** weights were plastic, updated according to Equation 6, whereas **W** and **Q** were held fixed throughout learning.

The networks contained four hidden areas of 500 units each and an output area of 10 units. Neurons in all areas used a sigmoid activation function and had a baseline burst probability of 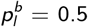. The **W, Y**, and **Q** weights were independently initialised with a Xavier normal initialisation. Networks were presented with MNIST images for 400 epochs using a batch size of 32, with the target class labels encoded as one-hot vectors at the output area. The burst-feedback weights **Y** were updated with a learning rate of *η*^(**Y**)^ = 0.04.

For experiments examining the effects of varying somatic noise, the number of apical branches, and teacher strength, the default settings were *σ*_noise_ = 0.4, *M*_*l*_ = 10 apical branches, and *κ* = 0.5. In each experiment, only one of these quantities was varied while the others were held fixed at these default values.

#### Separating excitatory and inhibitory inputs

To quantify excitatory and inhibitory feedback separately, we decomposed the event-feedback (**Q**) and burst-feedback (**Y**) weights into their positive and negative components. The excitatory and inhibitory apical inputs were defined as

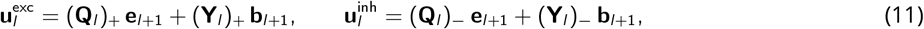

where (*x* )_+_ = max(*x*, 0) and (*x* )_−_ = min(*x*, 0) were applied element-wise, selecting the positive and negative weights, respectively. These components sum to the original apical potential, 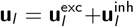, as defined in Equation 3. Excitatory and inhibitory inputs were compared across input examples before and after learning.

#### MNIST experiments

In these experiments, we evaluated whether plasticity of the burst-feedback weights, **Y**, could preserve **QY**-alignment during task learning. All experiments used the discrete-time rate-based BurstCCN, with **W** and **Y** plastic and **Q** fixed throughout training.

As in the previous set of experiments, the networks contained four hidden areas of 500 units each and an output area of 10 units, except where network depth was varied. Neurons in all areas used a sigmoid activation function and had a baseline burst probability of 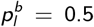. Networks were trained on MNIST images for 150 epochs using a batch size of 32, with target class labels encoded as one-hot vectors at the output area. Feedforward weights **W** were updated with a learning rate of *η*^(**W**)^ = 0.15 and weight decay of 5 *×* 10^−5^, while burst-feedback weights **Y** were updated with a learning rate of *η*^(**Y**)^ = 0.004. Independent somatic noise with standard deviation *σ*_noise_ = 0.1 was applied in all areas, the teaching signal strength was set to *κ* = 0.5, and, except in experiments where the number of apical branches was varied, the number of branches was *M*_*l*_ = 10.

For the online and hybrid BurstCCN models, training was organised into *online* and *realignment* blocks. During an online block, class labels were provided as teaching signals at the output area, and both feedforward weights **W** and burst-feedback weights **Y** were plastic. During a realignment block, teaching signals were removed, plasticity of **W** was disabled, and only **Y** remained plastic, allowing the burst-feedback pathway to realign with the fixed event-feedback pathway **Q** through the apical silencing rule in Equation 6. In the online regime, training consisted only of online blocks. In the hybrid regime, a realignment block was inserted after each training epoch.

For both the online and hybrid BurstCCN models, we initialised the network with imperfect **QY**-symmetry by perturbing the burst-feedback weights away from the **QY**-symmetric configuration. Specifically, for each area *l*,

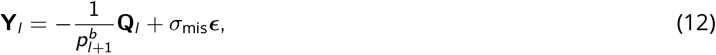

where 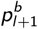 is the next area’s baseline burst probability, ***ϵ*** ∼ *N* (**0, I**), and *σ*_mis_ is the standard deviation of the perturbation. Here, *σ*_mis_ was set to 2% of the standard deviation of the initial **Q** weights. We additionally considered an ideal aligned regime, in which **Q** and **Y** were fixed in perfect **QY**-symmetry from the start of training such that no feedback realignment was required.

When comparing model types across network depth, the BurstCCN hybrid learning configuration used realignment blocks with a duration equal to 50% of the number of updates in the corresponding online blocks. The **QY**-symmetric BurstCCN networks were initialised without any misalignment between **Q** and **Y** weights and did not use **Y**-plasticity.

#### Representation analysis

To compare the representations learned by BurstCCN with those learned by artificial neural networks trained with feedback alignment, we analysed activity patterns in trained BurstCCN and ANN-FA networks. We first visualised just the second hidden-area representations of the online BurstCCN and ANN-FA using t-distributed stochastic neighbour embedding (t-SNE) ^58^, which projects high-dimensional activity patterns into a two-dimensional space. We then used representational trajectory analysis (RTA) ^35^ to compare how the representations of all models changed across successive network areas. For each model and area, this involved computing a representational dissimilarity matrix (RDM) from the corresponding activity vectors using Pearson correlations:

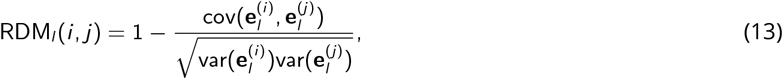

where 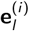 denotes the area-*l* activity vector for test example *i*, corresponding to event rates in BurstCCN and the feed-forward activities in ANN-FA. The off-diagonal upper-triangular entries of each RDM were vectorised to produce one representation vector per area. These vectors were then compared with each other by computing a meta-dissimilarity matrix (MDM), whose entries corresponded to the Pearson correlation distance between pairs of model-area representations. Embedding this MDM into two dimensions using multidimensional scaling (MDS) ^59^ produces a representational trajectory for each model through the network hierarchy. Models with similar trajectories therefore learned similar transformations of the input across areas, whereas divergent trajectories indicate qualitatively different learned representations.

#### CIFAR-10 and ImageNet experiments

For these experiments, we evaluated models on the CIFAR-10^60^ and ImageNet ^61^ datasets. We used the discrete-time convolutional BurstCCN (see Supplementary Methods), in which feedforward (**W**), burst-feedback (**Y**), and event-feedback (**Q**) weights were all plastic. Hidden areas used ReLU activation functions followed by divisive area normalisation, and the output area used a softmax function. The baseline burst probability was set to 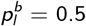 in all areas. The teaching-signal strength was set to *κ* = 0.1 for CIFAR-10 and *κ* = 0.001 for ImageNet. No somatic noise was applied in these experiments.

For CIFAR-10, we used a network consisting of three convolutional areas followed by a fully connected hidden area with 1500 units. For ImageNet, we used a deeper network with seven convolutional areas followed by a fully connected hidden area with 1500 units. Full architectural details are provided in Table S3. For both datasets, standard data augmentation and preprocessing were applied to the images during training.

We compared three feedback configurations. In the *random fixed* configuration, feedback weights were initialised randomly in a **QY**-symmetric configuration, fixed throughout training, and independent of the feedforward weights, analogous to Feedback Alignment (FA). In the *random plastic* configuration, feedback weights were also initialised randomly and independently of **W**, but were updated during training using the Kolen–Pollack (KP) plasticity rule. In the *symmetric plastic* configuration, feedback weights were initialised symmetrically with the feedforward pathway and updated using the same KP rule.

For CIFAR-10, BurstCCN models were trained with a batch size of 32, using learning rates of *η*^(**W**)^ = 0.002 and *η*^(**Y**)^ = *η*^(**Q**)^ = 0.001, with weight decay of 10^−4^ applied to all weight types. For ImageNet, BurstCCN models were trained with a batch size of 128, using learning rates of *η*^(**W**)^ = *η*^(**Y**)^ = *η*^(**Q**)^ = 0.01, again with weight decay of 10^−4^. For the baseline ANN models, we trained standard convolutional networks with the same architectures and hyperparameters, except for *η*^(**Q**)^, which applies only to the **Q** weights in BurstCCN.

#### Frozen Lake reinforcement learning task

We evaluated BurstCCN in a reinforcement-learning setting using the Frozen Lake navigation task ^62^. The agent navigated a 3 *×* 3 grid world and was trained to reach a fixed goal location while avoiding holes in the ice. Ice hole locations were randomly sampled and reshuffled every 10 episodes. At each timestep, the agent observed a binary state vector encoding both its current location and the hole locations. Specifically, the state consisted of two binary vectors of length 9, one for the agent position and one for the hole positions, giving an input dimension of 18. The agent selected one of four actions, corresponding to movement up, down, left, or right. A reward *R* = 1 was delivered upon reaching the goal, and *R* = 0 otherwise. Episodes terminated when the agent reached the goal or one of the ice hole locations.

We trained the agent using a deep Q-network (DQN) approach ^37^, with the action-value function implemented by a discrete-time rate-based BurstCCN. The network had 18 input units, three hidden areas of 64 units each, and four output units corresponding to the four possible actions. Each output unit represented the value of one action through its event rate. An experience replay buffer containing 20,000 timesteps was used, and a separate target network was updated from the policy network every 1,000 timesteps. The feedforward weights were trained with learning rate *η*^(**W**)^ = 0.5, with no momentum.

All areas used a sigmoid nonlinearity, which meant that output event rates were constrained to lie between 0 and 1. We therefore represented Q-values by rescaling them into the event-rate range:

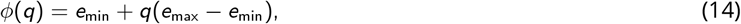

with *e*_min_ = 1*/*5 and *e*_max_ = 4*/*5, so that Q-values in [0, 1] were represented by output event rates in [*e*_min_, *e*_max_].

For a transition (*s, a, R, s*^*′*^), the target event rate for the selected action was computed as

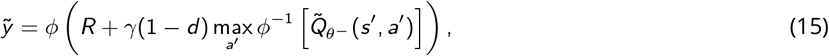

where *d* = 1 for terminal transitions and *d* = 0 otherwise, *θ*^−^ denotes the target-network parameters, 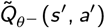 is the target network’s output event rate for action *a*^*′*^, and *ϕ*^−1^ denotes the inverse mapping from event-rate space back to Q-value space. The resulting 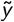 was used as the BurstCCN teaching signal for the selected output unit, while the targets for the unselected actions were left unchanged.

To assess whether trained BurstCCN activity represented task-relevant information during reinforcement learning, we recorded hidden-area activity as the agent interacted with the Frozen Lake environment. Linear decoders were trained to predict value-related variables, including action-value prediction error, selected action value and distance to the goal, as well as task-state variables, including the selected action, agent location and ice-hole locations. Continuous variables were decoded using ridge regression and categorical variables using logistic regression, with performance estimated by cross-validation.

### Model simulations of experimental observations

#### Interneuron silencing experiments

To compare the Dalean BurstCCN with experimentally observed interneuron-suppression effects on synaptic plasticity, we extracted paired pre- and post-stimulation L4-evoked postsynaptic potential amplitudes from Fig. 1g and Fig. 5c,g,j of Williams and Holtmaat ^38^ . For each paired recording, the relative change in synaptic strength was quantified as 100 *×* (post − pre)*/*pre. We then examined whether silencing analogous interneuron populations in the model produced corresponding changes in feedforward synaptic plasticity.

We used a Dalean BurstCCN consisting of two pyramidal neurons arranged in series. The first neuron received a constant input and projected to the second (output) neuron. For the output neuron, the excitatory pyramidal-to-pyramidal weights were initialised to 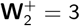, and the inhibitory PV-to-pyramidal weights were initialised to 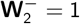, giving an effective net feedforward weight of **W**_2_ = 2. The bias for the output neuron was set to −0.5. For the first neuron, the feedforward weights were chosen such that the initial output event rate of the network was 0.5. The feedforward weights onto the first neuron were plastic and were updated according to the burst-dependent plasticity rule (Equations 5 and S18), while those onto the output neuron were held fixed to ensure that learning at the output neuron could not compensate for the effects of silencing each interneuron type. The feedback weights were initialised using the identity-based initialisation for the Dalean BurstCCN and were held fixed throughout learning.

To match the experimental condition, the task was set up to require LTP of the input weights by training the network to produce a target above its initial output event rate of 0.5. In each interneuron silencing condition, all interneurons of a given type (PV, SST, NDNF, or VIP) were completely silenced by setting their activity to 0, such that they provided no input to their postsynaptic targets. To generate variability across runs, the constant input and target output event rate were drawn independently for each run from the uniform distributions *e*_0_ ∼ *U* (0.8, 1.6) and *e*_target_ ∼ *U* (0.6, 1.0), respectively, and were held fixed throughout training. Networks were trained for 200 timesteps using a feedforward learning rate of *η*^(**W**)^ = 1.0. Synaptic plasticity was quantified as the total relative change in the input weights from their initial values, 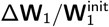, where 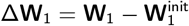.

#### Reduced SST population experiments

To test how the experimentally observed increase in SST interneuron density across the cortical hierarchy affects learning in the model, we varied the size of the SST interneuron population conveying burst-feedback in the Dalean BurstCCN. The SST density values used to show the cortical SST-density gradient were extracted from Fig. 2f of Kim et al. ^39^ . For these experiments, the network was trained on the Fashion-MNIST dataset ^63^ and consisted of four hidden areas of 50 pyramidal units each.

This experiment used a Dalean BurstCCN with the reduced-rank burst-feedback initialisation, in which the number of SST interneurons determines the effective dimensionality of the burst-feedback pathway. SST population size was defined as the total number of SST interneurons per area. At initialisation, the event-feedback weights, **Q**, were fixed in symmetry with the effective burst-feedback weights, **Y**, and both feedback pathways were held fixed throughout training. The feedforward weights, **W**^+^ and **W**^−^, were plastic and were updated according to their corresponding burst-dependent plasticity rules (Equations 5 and S18), using a learning rate of *η*^(**W**)^ = 0.15 and weight decay of 5 *×* 10^−5^.

We first varied the number of SST interneurons uniformly across areas. We then fixed the baseline population size at 8 SST interneurons per area and assessed the effect of reducing one area at a time to 4 SST interneurons. In each case, the apical feedback rank was quantified from the apical feedback activity matrix by centring activity across input examples, computing its singular values *s*_*i*_, and calculating the participation-ratio effective rank, 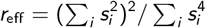. For the single-area reduced networks, this value was normalised to the corresponding baseline effective rank from a network with 8 SST interneurons in every area and expressed as a percentage.

#### Cue- and error-related responses in SST subpopulations

To compare the Dalean BurstCCN with experimentally observed SST subpopulation dynamics, we extracted SSTp and SSTn cue- and reward-response values from Fig. 7c of Chevy et al. ^41^ . Cue responses were calculated from fluorescence changes between cue onset and cue offset (−1.5 to −1s relative to reward delivery), whereas reward responses were calculated from fluorescence changes over the 1s following reward delivery (0 to 1s). We then examined whether analogous SST interneuron subpopulations in the model exhibited distinct stimulus- and error-related responses across learning. The network was trained on MNIST and contained four hidden areas of 500 units each with sigmoid activation functions. Feedback weights were initialised using the identity-based initialisation. All analyses were performed on the SST population in the second hidden area. For each input example, SST interneuron activity was measured first when the input image was presented without the target label and again when the input image was presented together with the target label. We defined the activity when the input image was presented without the target label as the stimulus response and the change in activity when the target label was presented as the error response. SST subpopulations were defined by ranking interneurons according to their error response before training, labelling the top 40% most responsive cells as SSTp and all remaining cells as SSTn. After training, stimulus and error responses were measured in the same way using the same SSTp and SSTn subsets. For each subpopulation, responses were averaged across neurons for each example, and reported values show the mean and standard error of these example-level responses.

#### Neuron-specific error signals in a BCI-inspired task

To examine whether BurstCCN can reproduce experimentally observed neuron-specific error signals, we implemented a simplified analogue of the BCI task setup used by Francioni et al. ^42^, in which the mean activity of two distinct neuronal populations controls the rotation of a visual grating. To recreate this task, we constructed a network consisting of 40 input units, two hidden areas of 40 pyramidal neurons each, and a single output neuron. Each input was drawn independently from a standard normal distribution. We then partitioned the first hidden area’s neurons randomly into two equal-sized populations of 20 neurons, denoted P+ and P−.

Analogous to the BCI setup, we defined a scalar control variable based on the difference between the mean activity of the two populations,

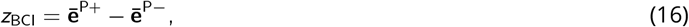

where 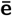 denotes the average event rate across neurons within each population. This quantity is an analogue of the grating angle in the experimental task.

In our setup, we first trained the network to predict this variable at the output area by providing a linearly rescaled version as the target for the sigmoid output unit,

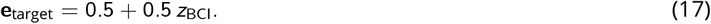

Training was performed for 5000 timesteps, during which the feedforward weights **W** between the two hidden areas and between the second hidden and output areas were updated according to the burst-dependent plasticity rule. The corresponding feedback weights **Y** and **Q** were kept fixed. This procedure establishes a readout in which P+ neurons contribute positively to the output and P− neurons contribute negatively, consistent with their opposing causal roles in the task. In the NDNF-activation condition, this training procedure was repeated with a constant input of 1 added to the NDNF interneuron activity throughout training.

We then examined the feedback signals received by hidden-area neurons under controlled target perturbations, with all weights held fixed. Rather than modelling the full closed-loop dynamics of the BCI task, we instead considered only the instantaneous instructive signal conveyed by the feedback pathways. We used two probe conditions in which the output-area target event rate was shifted relative to the network’s current output, 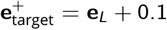 and 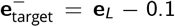. These perturbations provided analogues of the increasing- and decreasing-error regimes in the experimental setup. Neuron-specific error signals were quantified as the burst probability of each hidden-area pyramidal neuron, which reflects apical dendritic input in the model and serves as an analogue of the somato-dendritic residual measured experimentally. Baseline burst probability was set to 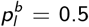. For each condition, burst probabilities were measured across 1000 input samples and five random initialisations. Distributions of *p*_*i*_ were computed separately for P+ and P− populations and visualised using empirical cumulative distribution functions.

The experimental reference distributions and summary values were obtained by extracting z-scored somato-dendritic residual values for P+ and P− neurons from Fig. 5e,g of Francioni et al. ^42^ .

## Code availability

The code for all implementations of BurstCCN is available at https://github.com/neuralml/BurstCCN-journal.

## Data availability

No new experimental datasets were generated in this study. The MNIST ^64^, Fashion-MNIST ^63^, CIFAR-10^60^, ImageNet ^61^ and CatCam ^24^ datasets are publicly available from their original sources. Previously published experimental data reproduced or adapted in Fig. 6 are available from the corresponding original publications.

## Supplementary Information

### Supplementary Figures

**Figure S1.**
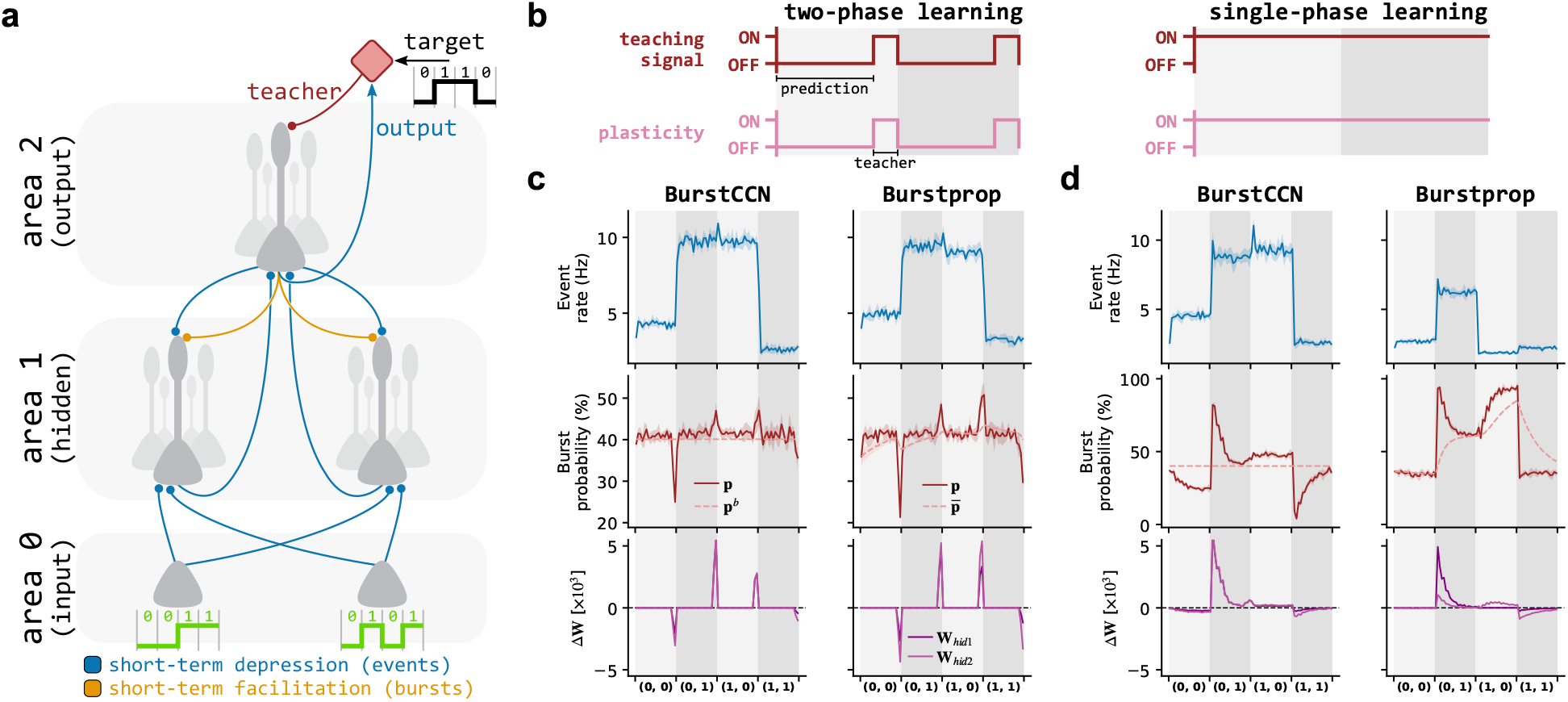
BurstCCN learns a nonlinear associative task without phased learning. **a**, Illustration of the hierarchical BurstCCN network trained on the nonlinear associative exclusive OR (XOR) task. **b**, Illustration of the two learning regimes. Left: Two-phase learning, with a prediction-only period (teaching signals and plasticity OFF) followed by a teaching period (teaching signals and plasticity ON; see Supplementary Methods). Right: Single-phase learning, where teaching signals and plasticity remain ON throughout. **c**,**d**, Learning dynamics under the two-phase (**c**) and single-phase (**d**) regimes. The left column shows BurstCCN and the right column shows Burstprop. *Top*: Output area event rates. *Middle*: Burst probability (**p**), along with its reference value (either the fixed baseline **p**^*b*^ in BurstCCN or the moving average 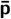 in Burstprop). *Bottom*: Weight updates of the hidden-to-output connections. Each line represents the mean, with standard error shown as shaded regions (*n* = 5).

**Figure S2.**
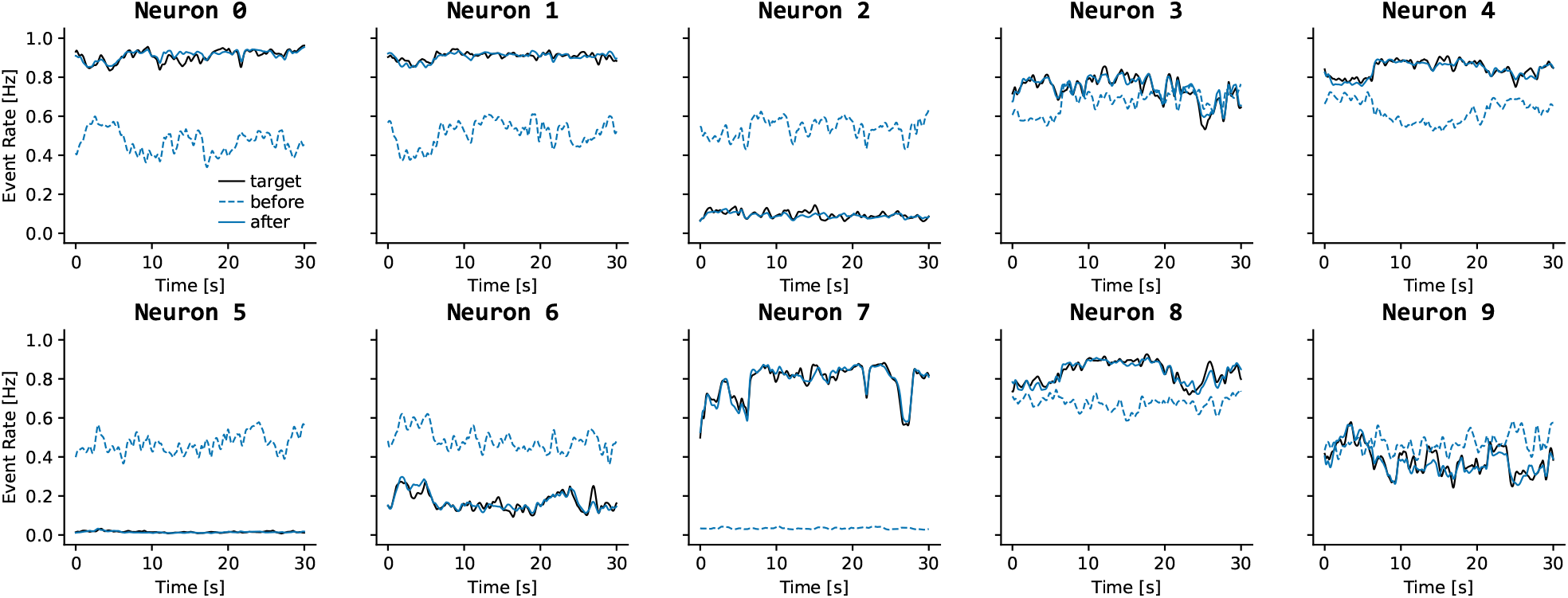
BurstCCN output neurons match their targets in the naturalistic regression task. All 10 output neurons (blue lines) learn to match their respective targets (black lines) for the naturalistic CatCam task.

**Figure S3.**
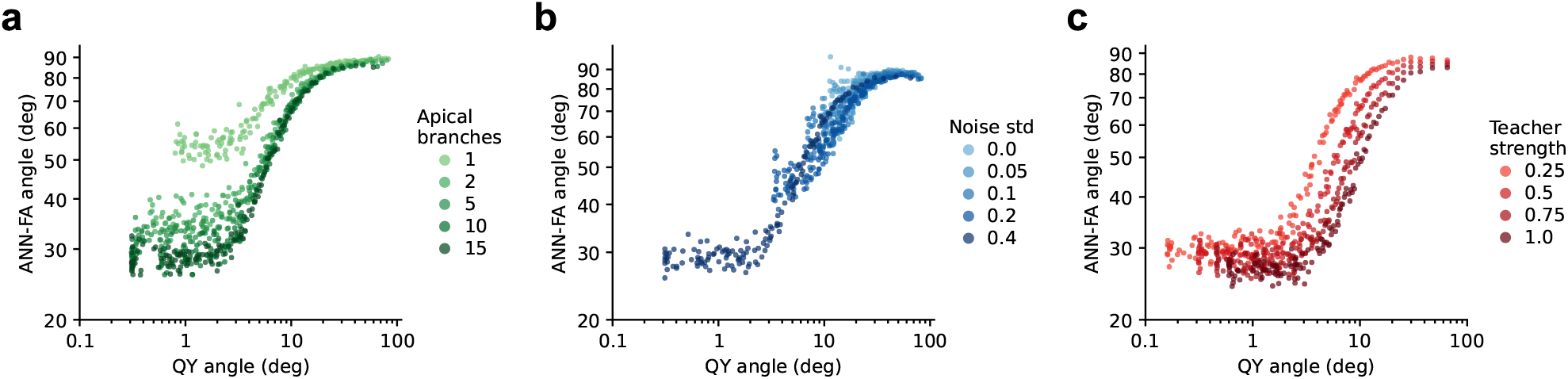
Feedback-pathway alignment is positively related to update alignment. Relationship between the **QY**-alignment angle and the ANN-FA update alignment angle over the course of **Y** plasticity for the model configurations shown in Fig. 3, in which the following factors were varied: **a**, the magnitude of independent somatic noise injected into downstream neurons; **b**, the number of apical branches receiving distinct **Q** and **Y** feedback inputs; and **c**, the teaching-signal strength to the output apical dendrites. Each point corresponds to the pair of alignment angles measured at a single timestep during **Y** plasticity. For visualisation, points were subsampled evenly across the range of **QY**-alignment angles.

**Figure S4.**
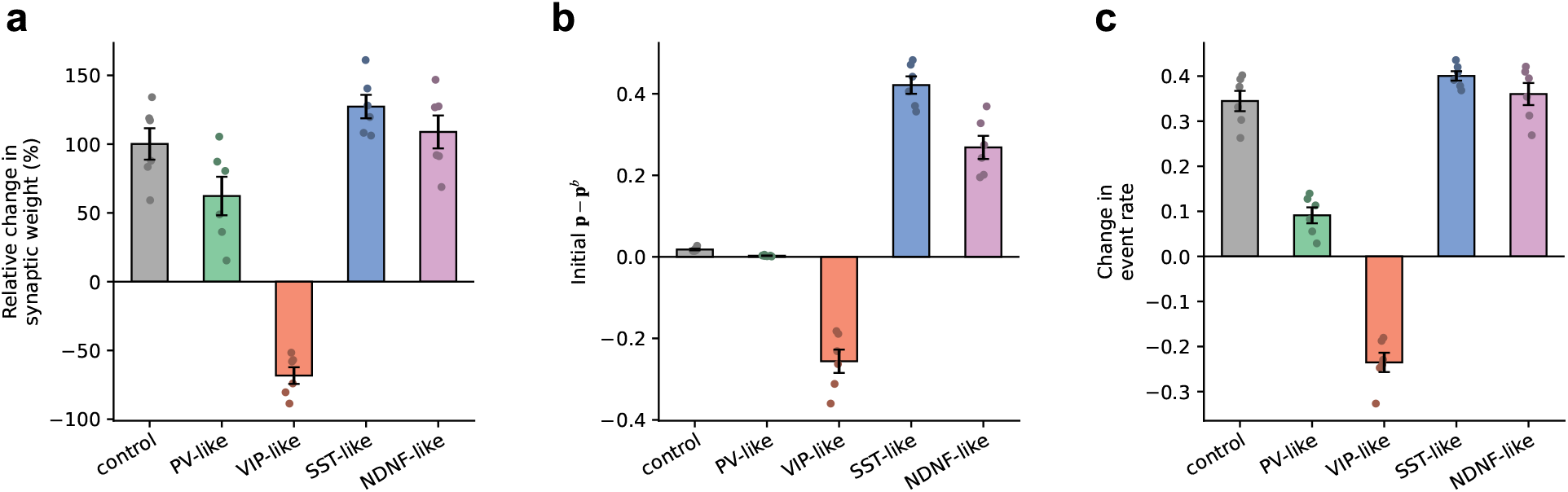
Additional cell-type silencing metrics for the Dalean BurstCCN. **a**, Relative changes in feedforward synaptic weights in the Dalean BurstCCN when individual interneuron populations were selectively silenced, corresponding to Fig. 6b, right. Changes are quantified as 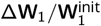. **b**, Initial change in burst probability relative to baseline for each cell-type silencing condition, measured at the start of silencing. **c**, Change in event rate from the start of silencing to the end of learning for each cell-type silencing condition.

**Figure S5.**
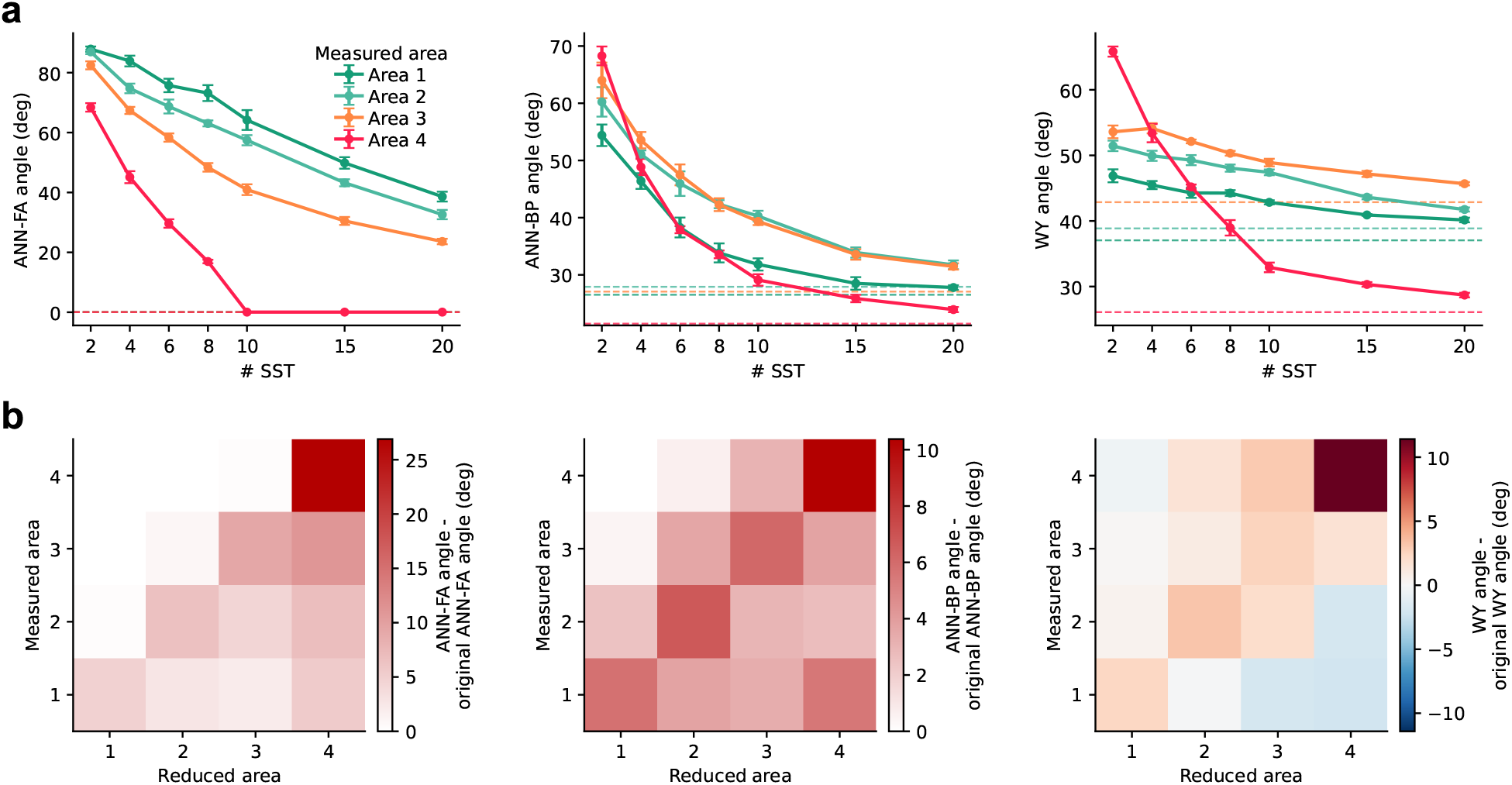
Alignment effects of SST population size in the Dalean BurstCCN. Effects of SST population size on update and weight alignment in the Dalean BurstCCN. **a**, Uniform changes in SST population size across all areas. Left: angle between Dalean BurstCCN updates and ANN-FA updates. Middle: angle between Dalean BurstCCN updates and ANN-BP updates. Right: angle between the feedforward weights **W** and burst-feedback weights **Y**. Dashed lines indicate the corresponding values with the full SST population size. **b**, Area-specific reductions in SST population size. Each heatmap shows the change in the corresponding angle relative to the same measured area with the full SST population size. Positive values indicate an increase in angle and therefore a loss of alignment, whereas negative values indicate a decrease in angle and therefore improved alignment.

**Figure S6.**
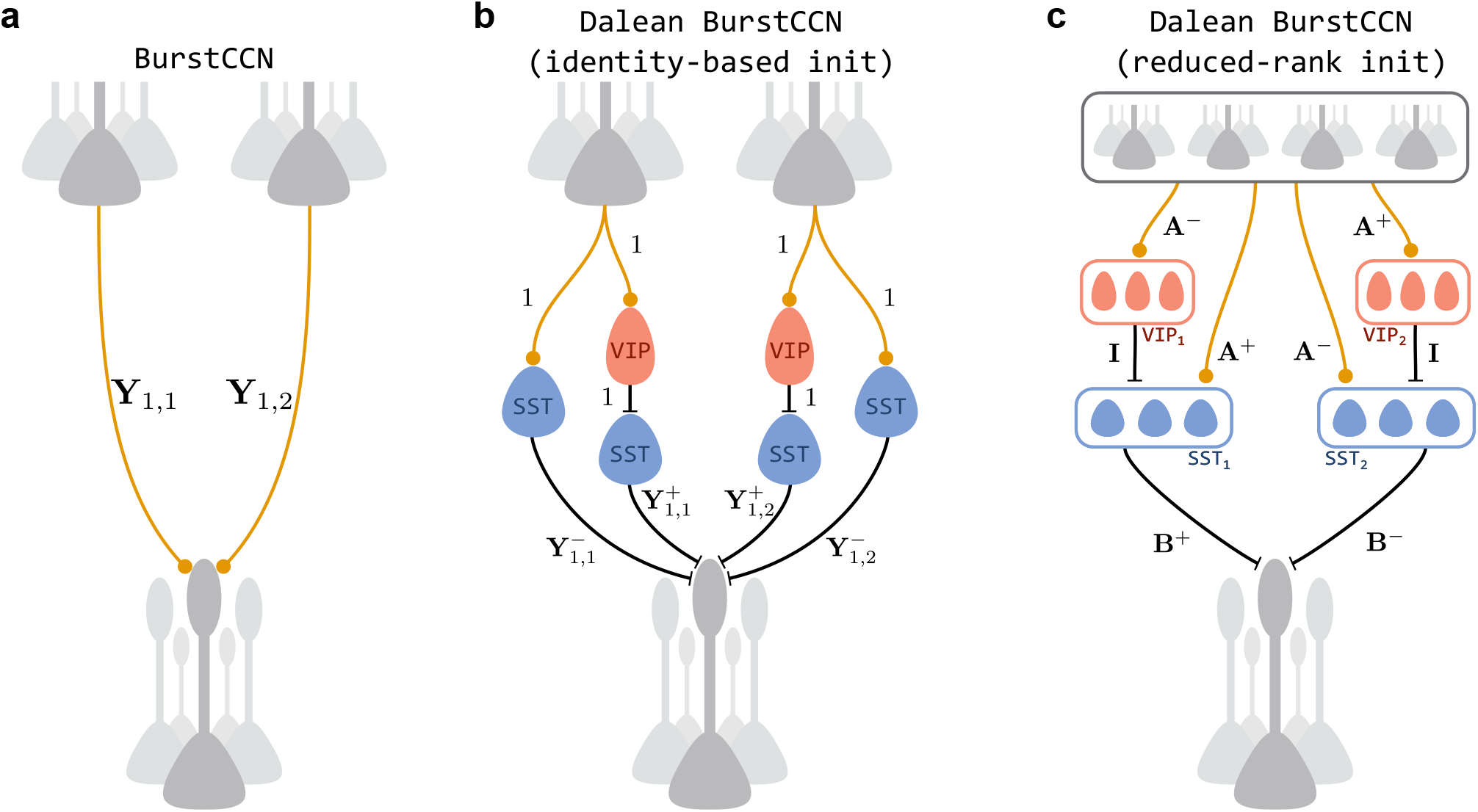
Burst-feedback pathway initialisations in the Dalean BurstCCN. Burst-feedback pathway weight initialisations in the Dalean BurstCCN, compared with the standard BurstCCN. **a**, Standard BurstCCN burst-feedback pathway. **b**, Identity-based Dalean initialisation, in which SST subpopulations carry complementary one-to-one representations of the next-area burst rates. **c**, Reduced-rank Dalean initialisation, in which burst feedback is implemented through a smaller SST/VIP population using a rank-constrained factorisation. Where shown, the subscripts on each labelled burst-feedback connection denote the target and source pyramidal units associated with that connection, respectively.

### Supplementary Methods

#### Spiking BurstCCN

This section describes the spiking BurstCCN which follows a similar approach to the spiking implementation of Burstprop ^11^, using the same two-compartment pyramidal neuron model developed by Naud and Sprekeler ^22^. This neuron model uses biophysically realistic dynamics in the somatic and apical dendritic compartments, which interact to generate patterns of single spikes and bursts that encode the inputs into each compartment. The spiking BurstCCN organises these neurons into ensembles, which themselves are organised into areas with the same ensemble-to-ensemble connectivity and short-term plasticity characteristics as the rate-based models. Unlike the rate-based models, which explicitly model the multiplexed burst ensemble code, in this spiking model, the burst multiplexing emerges from the biologically realistic neuron dynamics, providing a more accurate representation of multiplexed error communication across the network areas.

To keep the notation simple, this section considers the case where each area of the spiking BurstCCN consists of a single ensemble of *N* neurons. Neurons in the area *l* − 1 ensemble connect via feedforward synaptic weights, **W**_(*l*)_, to the somatic compartments of neurons in the area *l* ensemble. Neurons in the area *l* + 1 ensemble connect via two types of feedback synaptic weights, **Y**_(*l*)_ and **Q**_(*l*)_, onto the apical compartments of neurons in the area *l* ensemble. Note that in each case, these weight matrices are *N*-by-*N*, representing the connections between individual neurons in each ensemble. When this is extended to the case of multiple ensembles per area, these synaptic weight matrices exist between all pairs of ensembles in adjacent areas.

Each neuron, indexed by *i*, has two compartment potentials: a somatic potential, 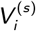, and an apical dendritic potential, 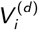. The dynamics of these potentials generate a pattern of activity characterised by a spike train, *S*_*i*_ (*t*), an event train, *E*_*i*_ (*t*), and a burst train, *B*_*i*_ (*t*), each defined as a binary function over time, *t*. These functions evaluate to 1 when the corresponding type of spiking event occurs and 0 otherwise. A spike occurs whenever the neuron exceeds its spike threshold. An event is defined as a spike that occurs after the neuron has not spiked for at least 16 ms. Following each event, if the neuron spikes again within 16 ms, this second spike is defined as a burst. Any subsequent spikes that occur before the next event are considered part of that burst.

#### Somatic dynamics

Events occurring in one area propagate through the **W**-weights to produce an input current in the soma of each neuron in the next area:

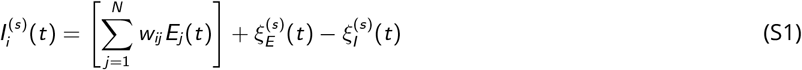

where *w*_*ij*_ is the feedforward synaptic weight between presynaptic neuron *j* in area *l* − 1 and postsynaptic neuron *i* in area *l*, corresponding to an element of **W**_(*l*)_. The terms 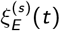 and 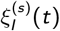 represent somatic excitatory and inhibitory noise inputs, each generated from independent Poisson processes with the same rate *λ*^(*s*)^ and amplitude *A*^(*s*)^. This noise plays an important role in linearising the response of neurons to their inputs and in decorrelating the spiking activity within each ensemble, both of which greatly improve the ability to decode the bottom-up somatic input from the ensemble activity ^22^.

The somatic input current directly influences the somatic potential of the neuron, 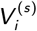, which evolves over time following:

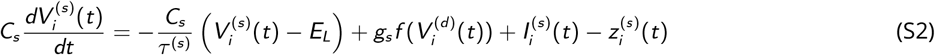

where *C*_*s*_ is the somatic capacitance, *τ* ^(*s*)^ is the somatic membrane time constant and *E*_*L*_ is the leak reversal potential. The term involving the function *f* introduces coupling with the dendritic compartment defined using a sigmoid function:

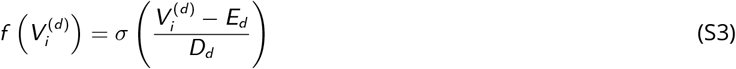

where *E*_*d*_ controls the midpoint of the sigmoid and *D*_*d*_ controls its slope. This nonlinear coupling, modulated by *g*_*s*_, provides a mechanism for the dendritic compartment potential to strongly drive depolarisation in the soma to induce bursting. The term 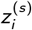 is a somatic spike-triggered adaptation variable that follows its own dynamics given by:

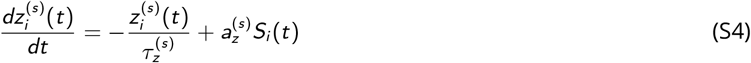

where 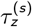 is the adaptation time constant, 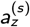 is the adaptation strength and *S*_*i*_ (*t*) corresponds to the spike-train for this cell. This is a form of spike-frequency adaptation, regulating the neuron’s firing rate through a feedback mechanism that reduces its excitability as firing increases. Its effect is to smooth each neuron’s response, further promoting a more linear input-output relationship.

A spike is generated by the neuron when the somatic potential exceeds a dynamic spike threshold, 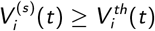, after which the somatic potential is instantaneously reset back to a constant voltage, *V*_*reset*_ . This threshold evolves over time according to:

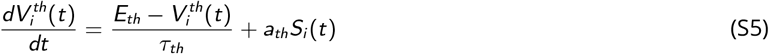

where *τ*_*th*_ is the threshold time constant, *E*_*th*_ is the threshold reversal potential and *a*_*th*_ controls the increase in threshold following each spike. In contrast to the spike-frequency adaptation, the spike-threshold adaptation operates on a shorter timescale and functions as a refractory mechanism that prevents many spikes in quick succession.

#### Dendritic dynamics

The apical dendritic input current differs between neurons in the output area and those in the hidden areas. For neurons in the output area:

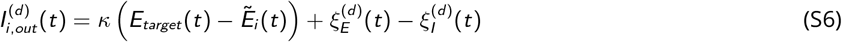

where *E*_*target*_ (*t*) is the target event rate, 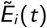 represents an exponential moving average of the event rate *E*_*i*_ (*t*), and *κ* controls the strength of the teaching input. The terms 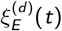 and 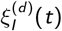 denote dendritic excitatory and inhibitory noise inputs, each generated from independent Poisson processes with the same rate *λ*^(*d*)^ and amplitude *A*^(*d*)^. Similar to the somatic noise, these terms play an important role in linearising the dendritic response to inputs and in decorrelating the dendritic activity across the ensemble, both of which greatly improve the population-level representation and decodability of the top-down inputs.

For neurons in the hidden areas, the dendritic input is given by:

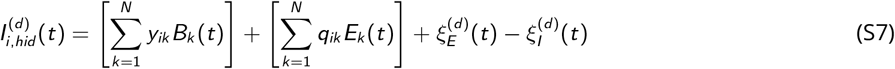

where *y*_*ik*_ and *q*_*ik*_ are the feedback synaptic weights from presynaptic neuron *k* in area *l* + 1 to postsynaptic neuron *i* in area *l*, corresponding to elements in **Y**_(*l*)_ and **Q**_(*l*)_, respectively. Note that the *y* weights (STF) transmit burst events and the *q* weights (STD) transmit all events corresponding to their assumed STP dynamics.

The dendritic input current directly influences the neuron’s apical dendritic potential, 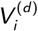, which evolves over time following nonlinear dynamics:

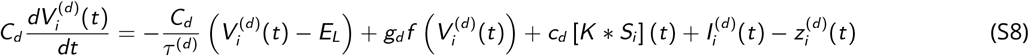

where *C*_*d*_ is the dendritic capacitance and *τ* ^(*d*)^ is the dendritic membrane time constant. Regenerative dendritic activity, modulated by *g*_*d*_, is modelled using the same voltage-dependent sigmoidal nonlinearity, *f*, defined in Equation S3. When the dendritic potential reaches a critical level of depolarisation, the nonlinear effect of this regenerative activity causes it to become unstable, amplifying the depolarisation in a self-reinforcing manner to produce a dendritic spike. Due to the dendritic compartment’s influence on the soma, triggering this threshold-like mechanism switches the neuron into an active bursting state. Additionally, the term [*K* * *S* ] (*t*) models the effect of backpropagating action potentials by convolving the spike train with a box-filter kernel, *K* (*t*) = 1 if 0.5ms ≤ *t* ≤ 2.5ms and 0 otherwise, which spreads the effect of each spike over this brief time window. The strength of these backpropagating action potentials is controlled by the scaling factor *c*_*d*_ . Including this mechanism is important for the burst code as it allows somatically induced spikes to conditionally trigger bursts, depending on the level of apical dendritic depolarisation. A single event occurs in either case, but the likelihood of triggering a burst increases with the dendritic potential. Lastly, the term 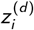 is a subthreshold dendritic adaptation variable that evolves according to:

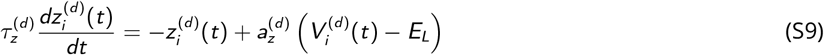

where 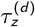 is the dendritic adaptation time constant and 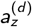 controls the strength of the adaptation. The role of this adaptation is to reset the dendritic potential back towards its reversal potential after a dendritic spike, preventing it from inducing long bursts of activity. It operates on a slower timescale than the decay governed by *τ* ^(*d*)^ in Equation S8, producing a more sustained current to overcome the effects of regenerative dendritic activity.

For simplicity, the spiking BurstCCN does not implement plasticity rules for **Q** and **Y** weights and instead assumes them to be fixed in the **QY**-symmetric state. Specifically, this means that 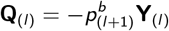, where 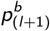 is the baseline burst probability for neurons in the area *l* + 1 ensemble. Additionally, unless otherwise stated, the hyperparameters of the spiking model in each experiment were set as shown in Table S1.

**Table S1.** Spiking BurstCCN default hyperparameters. The default set of hyperparameters used for experiments involving the spiking BurstCCN implementation, equal to the equivalent parameters used with the spiking Burstprop implementation ^11^.

| Parameter | Notation | Value |
| --- | --- | --- |
| Leak reversal potential | $E_L$ | -70mV |
| Somatic membrane time constant | $\tau^{(s)}$ | 16ms |
| Dendritic membrane time constant | $\tau^{(d)}$ | 7ms |
| Somatic capacitance | $C_s$ | 370pF |
| Dendritic capacitance | $C_d$ | 170pF |
| Somatic conductance | $g_s$ | 1300pA |
| Dendritic conductance | $g_d$ | 1200pA |
| Dendritic activation threshold | $E_d$ | -38mV |
| Dendritic activation slope | $D_d$ | 6mV |
| Backpropagating action potential scaling factor | $c_d$ | 2600pA |
| Somatic adaptation time constant | $\tau_z^{(s)}$ | 100ms |
| Dendritic adaptation time constant | $\tau_z^{(d)}$ | 30ms |
| Somatic adaptation strength | $a_z^{(s)}$ | 200pA |
| Dendritic adaptation strength | $a_z^{(d)}$ | 13nS |
| Somatic reset potential | $V_{reset}$ | -70mV |
| Spike threshold reversal potential | $E_{th}$ | -50mV |
| Spike threshold time constant | $\tau_{th}$ | 27ms |
| Spike threshold adaptation strength | $a_{th}$ | 2mV |

**Table S2.**
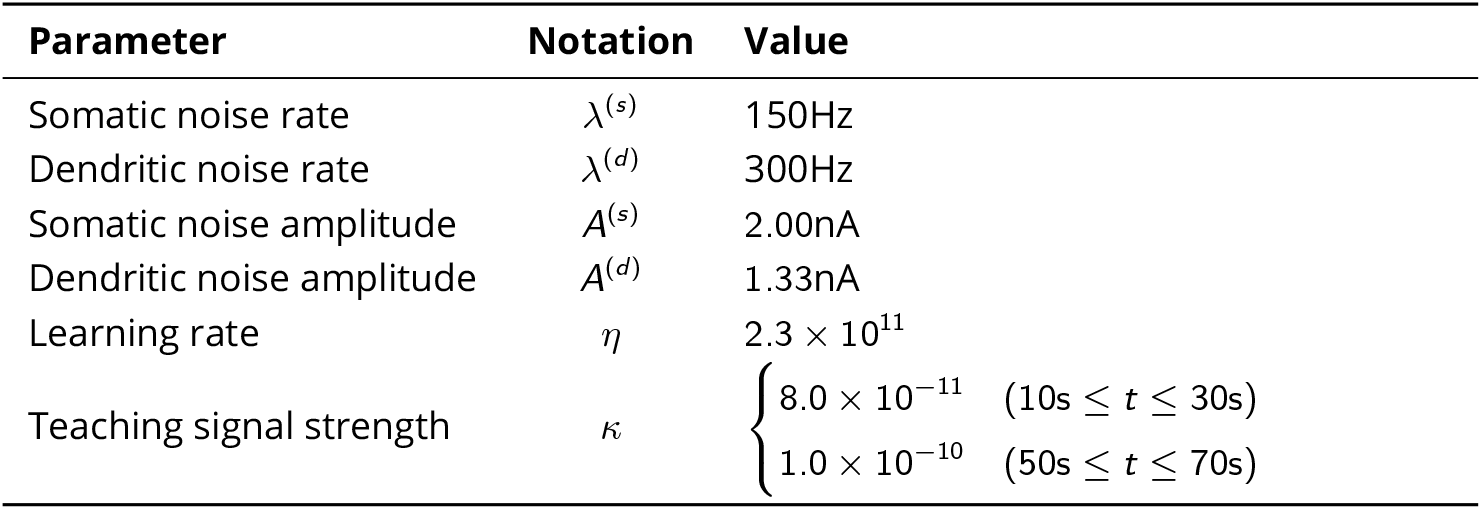
Spiking BurstCCN hyperparameters for the stationary target prediction task. The characteristics of somatic and dendritic noise and learning rates, used throughout both task stages, and the task-stage-specific teaching signal strengths.

#### XOR task

For this task, the network consisted of five neuron ensembles: two input ensembles, one for each input, two hidden ensembles, and one output ensemble. Each ensemble contained 500 individual neurons, resulting in a total of 2500 pyramidal neurons in each network simulation. Ensemble connectivity followed the same structure as in the task described above, and all weight matrices were sparse, with a connection probability of 5%. For both models, the burst feedback weights (**Y**) were fixed and initialised such that all connections were excitatory onto one of the hidden area populations and inhibitory onto the other. In the case of the BurstCCN, the additional event feedback weights (**Q**) were fixed in the **QY**-symmetric regime and the baseline burst probabilities were set to 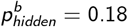 and 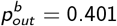 in the hidden and output areas, respectively. The two-phase setups used a feedforward learning rate of *η* = 4 *×* 10^−3^ (and *η*_*B*_ = 10^−4^ for weights from a set of bias units) which was reduced in the single-phase setups to *η* = 4 *×* 10^−4^ (and *η*_*B*_ = 10^−5^) to account for the increased duration of plasticity. These examples were repeatedly presented to the network a total of 2,000 times each in a deterministic order. The inputs were encoded as positive (200pA) and negative (−200pA) currents, corresponding to 1 and 0, respectively. Each example was presented for a total of 8s before the next example was shown, giving a total training time of 16,000s.

The two-phase learning regime has an initial prediction phase, lasting 7.2s for each input presentation. This is followed by a teacher phase for the remaining 0.8s during which plasticity is restored and teaching signals are provided at the output. In contrast, the single-phase regime does not involve a prediction phase at all and instead extends the teacher phase to the full duration of the input stimulus (Fig. S1b).

#### Continuous-time rate-based BurstCCN

In contrast to the discrete-time implementation, the continuous-time rate-based BurstCCN does not process each input example independently using just a single timestep and does not involve parallel processing using mini-batches. Instead, the network dynamics evolve over many continuous timesteps and are influenced by the history of the prior network states. The dataset of discrete input-output pairs is replaced by time-varying functions defining the sensory inputs, *x* (*t*), and output targets, *y* (*t*).

For each timestep, the input area event rates are instantaneously set to **e**_0_(*t*) = *x* (*t*) and the somatic potentials evolve following:

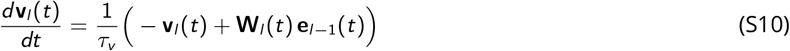

where *τ*_*v*_ represents the membrane leak time constant of the soma. Event rates are computed as an instantaneous function of these somatic potentials, **e**_*l*_ (*t*) = *f* **v**_*l*_ (*t*) . At the output area, burst probabilities are computed as:

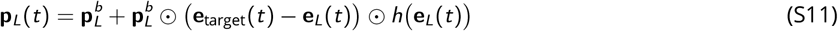

where, **e**_target_(*t*) = *y* (*t*), are the output-area targets and *h* is defined the same as in Equation 2. Note that the baseline burst probability, 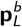, is constant and not a function of time. Burst rates are then computed as **b**_*l*_ (*t*) = **p**_*l*_ (*t*) ⊙ **e**_*l*_ (*t*), influencing the hidden area apical dendritic potentials that evolve with:

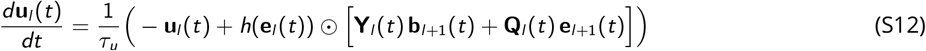

where *τ*_*u*_ is the membrane leak time constant of the apical dendrites. This results in the hidden area burst probabilities, 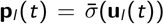, which further backpropagate through the areas of the network within the burst rates. As in the discrete-time model, the change in the burst probabilities of each area represents the assigned credit used for learning. The feedforward weights update over time following:

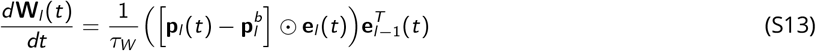

where *τ*_*W*_ is a synaptic time constant controlling the learning rate.

#### Convolutional architectures for CIFAR-10 and ImageNet experiments

For the complex image classification experiments, we replaced fully connected hidden areas with convolutional areas. In these areas, somatic potentials were computed by a convolution of the input feature maps with feedforward weights, **W**. Thus, for area *l*,

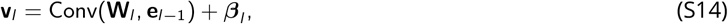

where ***β***_*l*_ denotes a bias term, and event rates were given by **e**_*l*_ = *f* (**v**_*l*_ ), using the same nonlinearity as in the fully connected model.

Feedback inputs to the apical compartment were computed using transposed convolution operations with separate burst- and event-feedback weights, **Y** and **Q**, which projected activity from area *l* + 1 back onto area *l* . The apical potentials were given by

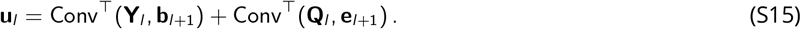

Burst probabilities and burst rates were computed as in the discrete-time fully connected model, with the relevant functions applied element-wise across feature maps. Feedforward convolutional weights were updated using the same burst-dependent plasticity rule as in Equation 5, with the outer product replaced by the standard convolutional weight update over local receptive fields.

#### ReLU activations with divisive area normalisation

For the complex image classification experiments, we used ReLU activation functions, *f* (**v**_*l*_ ) = ReLU(**v**_*l*_ ), followed by divisive normalisation across each area, implemented using an RMS-based normalisation rule ^65^. From the somatic potential **v**_*l*_, unnormalised event rates were computed as 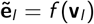 and were then normalised according to 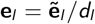, where the area-wise divisive scalar was

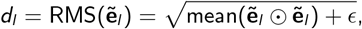

and *ϵ* is a small positive constant for numerical stability.

This normalisation couples the feedforward event rates of neurons within the same area. As a result, the credit assigned to each neuron is also no longer independent, but it depends on the credit assigned to all other neurons in the area. For RMS-normalised ReLU areas, the apical potential becomes

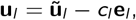

where **ũ**_*l*_ is the uncorrected apical potential computed as before using Equation 3, and

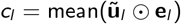

is an area-wise scalar correction term. This correction term can be approximated as 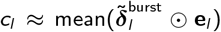, where 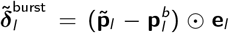 denotes the uncorrected difference in burst rate from baseline, computed using the apical potential **ũ**_*l*_, before subtracting the RMS-normalisation correction term. In other words, both *d*_*l*_ and *c*_*l*_ are area-wise scalar terms that can be expressed solely in terms of event and burst rates, quantities that are decodable across the area, without requiring access to subcellular compartment potentials.

**Table S3.** Convolutional architectures used for CIFAR-10 and ImageNet. Convolutional architectures used for both ANN and BurstCCN models in the CIFAR-10 and ImageNet experiments. CIFAR-10 inputs were 32 *×* 32 *×* 3 images, and ImageNet inputs were 224 *×* 224 *×* 3 preprocessed images. For convolutional areas, the table reports the number of output channels, kernel size, and stride. For fully connected areas, it reports the number of units.

| Task | Area | Area type | Channels/units | Kernel size | Stride |
| --- | --- | --- | --- | --- | --- |
| CIFAR-10 | 1 | Convolutional | 64 | $5 \times 5$ | 2 |
| | 2 | Convolutional | 128 | $5 \times 5$ | 2 |
| | 3 | Convolutional | 256 | $3 \times 3$ | 1 |
|  | 4 | Fully connected | 1500 | – | – |
|  | 5 | Fully connected | 10 | – | – |
| ImageNet | 1 | Convolutional | 48 | $9 \times 9$ | 4 |
| | 2 | Convolutional | 96 | $3 \times 3$ | 2 |
| | 3 | Convolutional | 96 | $5 \times 5$ | 1 |
| | 4 | Convolutional | 192 | $3 \times 3$ | 2 |
| | 5 | Convolutional | 192 | $3 \times 3$ | 1 |
| | 6 | Convolutional | 384 | $3 \times 3$ | 2 |
| | 7 | Convolutional | 470 | $3 \times 3$ | 1 |
|  | 8 | Fully connected | 1500 | – | – |
|  | 9 | Fully connected | 1000 | – | – |

#### BurstCCN with Dale’s Law constraints

Inspired by Dale’s ANNs ^66^, we introduce Dale’s law constraints to the BurstCCN by adding intermediate inhibitory interneuron populations between each area of pyramidal neurons. In contrast to Dale’s ANNs, which constrain only the feedforward connections, the Dalean BurstCCN also constrains the two feedback pathways. The schematic in Figure 6a shows how each connection type in the standard BurstCCN is implemented using excitatory and inhibitory pathways mediated by interneuron populations. Importantly, these constraints only change how feedforward and feedback signals are transmitted between areas. The local pyramidal-cell computations remain unchanged from the standard discrete-time BurstCCN: somatic and apical inputs determine compartmental potentials, which in turn determine event rates, burst probabilities, and burst rates. For simplicity, all interneuron populations are modelled as linear rate units.

The feedforward weights, **W**, are split into a direct excitatory pathway and an indirect inhibitory pathway, both targeting the somatic compartment of the next-area pyramidal cells. We interpret the inhibitory interneurons in this pathway as PV interneurons, consistent with their soma-targeting connectivity and short-term depressing dynamics ^44,46^. We denote **W**^+^ as the direct excitatory weights from pyramidal cells in area *l* − 1 to pyramidal cells in area *l*, 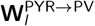 as the excitatory weights from pyramidal cells in area *l* − 1 to PV interneurons, and 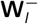 as the inhibitory PV-to-pyramidal weights in area *l* . The PV activity is given by

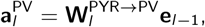

and the resulting somatic potential of pyramidal cells in area *l* is

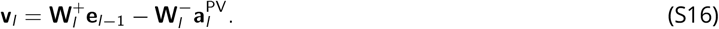

The event-feedback pathway (**Q**) is similarly split into a direct and an indirect pathway, both targeting the apical dendrites of the previous-area pyramidal cells. We interpret the inhibitory interneurons in this pathway as NDNF interneurons, consistent with their apical-targeting connectivity and short-term depressing dynamics ^67^. We denote **Q**^+^ as the direct pyramidal-to-pyramidal excitatory event-feedback weights, 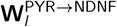 as the excitatory weights from pyramidal cells in area *l* + 1 to NDNF interneurons, and 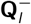 as the inhibitory weights from NDNF interneurons onto pyramidal apical dendrites in area *l* . The NDNF population activity is given by

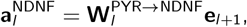

which gives the total event-related apical input

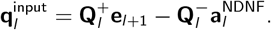

The burst-feedback pathway (**Y**) is not split in the same way to avoid introducing short-term facilitating pyramidal-to-pyramidal feedback onto apical dendrites, consistent with experimental observations that these connections exhibit short-term depressing dynamics ^68^. Instead, we implement burst-feedback through apical-targeting SST interneurons that receive inhibitory input from VIP interneurons to produce apical disinhibition. For notational simplicity, we write this SST/VIP-mediated feedback circuit using two complementary VIP and SST subpopulations, which should be understood as partitions of each interneuron population rather than as distinct biological interneuron classes with different circuit functions. We denote 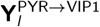 and 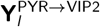 as the excitatory burst-feedback weights from pyramidal cells in area *l* + 1 to the two VIP subpopulations, 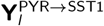 and 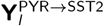 as the excitatory burst-feedback weights to the two SST subpopulations, 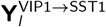 and 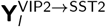 as the inhibitory weights from each VIP subpopulation to its corresponding SST subpopulation, and 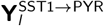 and 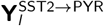 as the inhibitory weights from the two SST subpopulations to the apical dendrites of pyramidal cells in area *l* .

The VIP activity is determined by the burst rates of pyramidal cells in the next area:

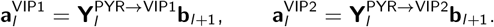

The SST interneurons depend on the same burst rates, together with inhibitory input from the matched VIP subpopulation:

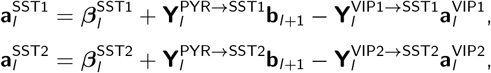

where 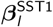 and 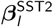 are excitatory bias terms. These biases allow the SST interneurons, which are constrained to be non-negative rate units, to support both increases and decreases in their inhibitory output onto pyramidal apical dendrites. The resulting burst-related apical input is then given by the combined SST-mediated pathways:

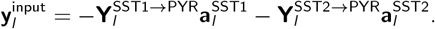

The total apical potential of pyramidal cells is obtained by combining the event-feedback and burst-feedback pathways with a bias term:

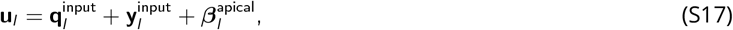

where 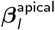 denotes an excitatory apical bias input.

We use two initialisations for the synaptic weights in this burst-feedback pathway, referred to as the identity-based and reduced-rank initialisations, which are described in detail below and illustrated in Fig. S6. In the identity-based initialisation, the SST1 and SST2 subpopulations carry complementary one-to-one representations of the next-area burst rates, allowing the circuit to implement full-rank burst feedback to the apical dendrites. In the reduced-rank initialisation, we use a singular value decomposition (SVD) of the burst-feedback weights **Y**_*l*_ to replace this one-to-one mapping with a smaller set of low-dimensional burst-rate projections, yielding a rank-constrained approximation of the original effective **Y**_*l*_ mapping.

The plasticity rules for the feedforward pathway are modified due to the presence of the intermediate PV interneuron population. The excitatory weights 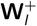 use the same update rule as in Equation 5, while the inhibitory weights 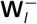 are updated according to:

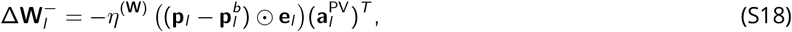

where the presynaptic activity now represents the PV interneuron activity. Throughout learning, all weights are constrained to remain non-negative by clipping negative values to zero immediately after each update.

#### Initialising feedforward and feedback pathways in the Dalean BurstCCN

Initialising the weights in the Dalean BurstCCN involved first initialising the corresponding signed weight matrices of the standard BurstCCN, **W**_*l*_, **Q**_*l*_, and **Y**_*l*_, across all areas. To obtain sign-constrained weights, we decomposed these signed matrices into their positive and negative parts. Specifically, we define the element-wise functions (*x* )_+_ = max(*x*, 0) and (*x* )_−_ = max(−*x*, 0), which extract the positive weights and the magnitudes of the negative weights, respectively. We denote these non-negative components using superscripts, such that 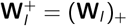 and 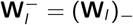, with analogous notation used for **Q**_*l*_, **Y**_*l*_, and any other signed matrices decomposed into positive and negative parts. This notation corresponds to the direct excitatory and indirect inhibitory weight matrices introduced in the Dalean BurstCCN formulation.

In all experiments, for the feedforward and event-feedback pathways, we initialise the pyramidal-to-interneuron weights as identity matrices:

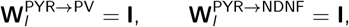

such that the PV and NDNF populations simply represent a copy of their presynaptic event rate inputs:

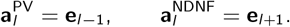

In the burst-feedback pathway, we consider two initialisation approaches: an identity-based initialisation, represented in Figure 6a, and a reduced-rank initialisation in which the SST subpopulations do not map one-to-one with pyramidal cells, allowing the number of SST cells to vary. In both cases, the SST bias terms are set according to

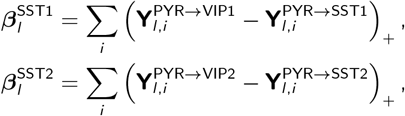

where **Y**_*l,i*_ denotes the vector of outgoing synaptic weights from presynaptic pyramidal neuron *i* to the corresponding interneuron population.

The apical bias term is then given by

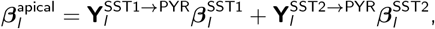

which precisely cancels the constant inhibition introduced by the SST bias terms.

#### Identity-based burst-feedback initialisation

In the identity-based initialisation, the burst-feedback circuit is initialised as

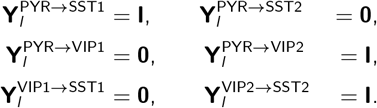

The weights from the two SST subpopulations to the pyramidal apical dendrites are then initialised as

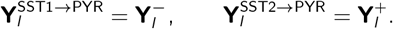

These two SST-to-pyramidal pathways therefore correspond directly to the negative and positive components of the effective burst-feedback matrix.

The VIP activities become

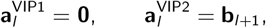

and the bias vectors reduce to

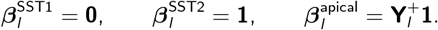

The SST activities then become

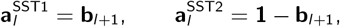

so that

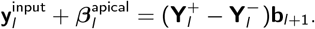

Hence the somatic and apical potentials simplify to

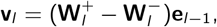

and

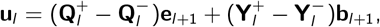

which recovers the original BurstCCN computation, since 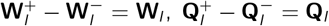, and 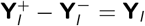.

#### Reduced-rank burst-feedback initialisation

To study the effect of limiting the dimensionality of burst-feedback communication, we reduced the size of the SST subpopulations in the burst-feedback pathway. Rather than allowing these SST subpopulations to copy the burst rates of the presynaptic pyramidal area with one-to-one identity connectivity, we approximated the original signed burst-feedback matrix **Y**_*l*_, of size 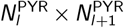, by a rank-constrained factorisation through *N*^SST^ SST cells. Specifically, we introduced a matrix **A**_*l*_, with dimensions 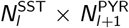, which maps burst rates in area *l* + 1 onto the SST and VIP populations, and a matrix **B**_*l*_, with dimensions 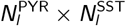, which maps SST activity onto the apical dendrites of pyramidal cells in area *l*. Reducing 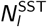 imposes a rank constraint on the burst-feedback pathway, thereby limiting the dimensionality of the feedback signal used for credit assignment.

To construct this reduced-dimensional mapping while preserving the original burst-feedback mapping as closely as possible, we used the singular value decomposition:

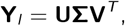

where **U** and **V** contain the left and right singular vectors of **Y**_*l*_, and **∑** is a diagonal matrix of singular values. For a reduced SST population of size 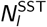, we retained the largest 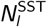 singular values and their corresponding singular vectors, giving the best rank-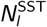 least-squares approximation of **Y**_*l*_ . Denoting these truncated matrices by 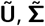, and 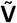, the two factor matrices were then defined as

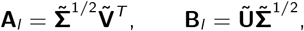

so that the approximate burst-feedback mapping becomes

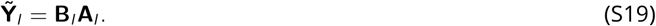

In the burst-feedback pathway, factor **A**_*l*_ was split element-wise into its positive and negative components, giving the weights to the SST and VIP subpopulations as

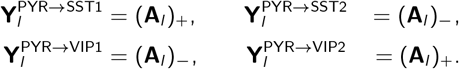

The VIP-to-SST weights were initialised as one-to-one inhibitory connections, with 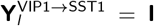 and 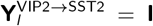. The factor **B**_*l*_ was then split as before into the two non-negative SST-to-pyramidal weight matrices

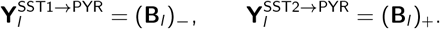

This factorised construction follows the same logic as the identity-based initialisation, but the positive and negative components are now defined at the level of the two matrix factors rather than by directly decomposing the full effective **Y**_*l*_ matrix. When 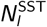 is at least as large as the rank of **Y**_*l*_, this construction can recover the original effective burst-feedback mapping exactly. For smaller SST and VIP subpopulations, it provides the closest rank-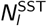 approximation of the original burst-feedback mapping.

#### Feedback learning leads to QY-symmetry

In this section, we demonstrate that the local update rule for the STF feedback weights, **Y**, leads to the **QY**-symmetric state necessary for the approximation to backprop to hold. The **QY**-symmetric state is defined as

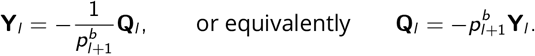

In particular, we show that in the absence of a teaching signal at the output area (i.e. **i**_teacher_ = 0) and assuming independently varying somatic potentials within the population, **Y**_*l*_ will converge to this solution.

First, we note that Equation 6 is simply implementing a gradient descent update rule with respect to the squared L2 norm of the apical potential:

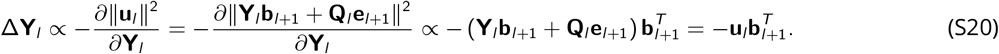

Following this gradient (with appropriate learning rate 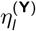 should ensure that a local minimum of ∥**u**_*l*_ ∥^2^ is reached. However, since **u**_*l*_ = **Y**_*l*_ **b**_*l*+1_ + **Q**_*l*_ **e**_*l*+1_ is linear in **Y**_*l*_, the objective ∥**u**_*l*_ ∥^2^ is quadratic in **Y**_*l*_ and therefore convex. Hence any local minimum is also a global minimum. In the absence of a teaching signal, the **QY**-symmetric solution enforces exact cancellation:

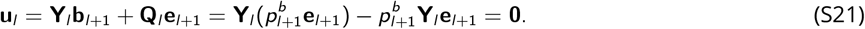

Thus the **QY**-symmetric solution achieves the global minimum where ∥**u**_*l*_ ∥^2^ = ∥**0**∥^2^ = 0. Moreover, in the case where burst activities **b**_*l*+1_ are linearly independent, it is a unique solution. Linear independence of **b**_*l*+1_ is not assured a priori, however, this can be guaranteed by injecting independent noisy currents into the population, as described in the Methods.

#### BurstCCN analytical mapping onto the backpropagation algorithm

In this section, we analytically show that, under mild assumptions, the feedback pathway of BurstCCN communicates signals that approximate the error gradients computed by backprop. We first outline how backprop defines error gradients recursively in terms of those in the next area, using notation consistent with BurstCCN. Let 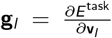 denote the error gradient with respect to the somatic potentials (i.e. pre-activations) at area *l*. Backprop involves computing **g**_*l*_ recursively with:

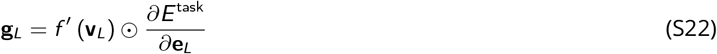

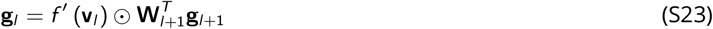

where *f* ^*′*^ denotes the activation function derivative of *f* . The error gradients with respect to the feedforward weights onto area *l* are then simply obtained as:

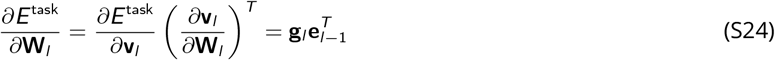

For BurstCCN, we consider the change in burst rate from its baseline level in each area, 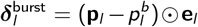, and show that this quantity mirrors −**g**_*l*_ in backprop. Note that this negative relationship arises from a choice of sign, such that increases in bursting correspond to synaptic potentiation (LTP), consistent with experimental observations ^15,69^. In the output area, by the definition of the burst probabilities (Equation 2), the change in burst rate 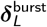 relates exactly to the backpropagated gradient **g**_*L*_ as follows:

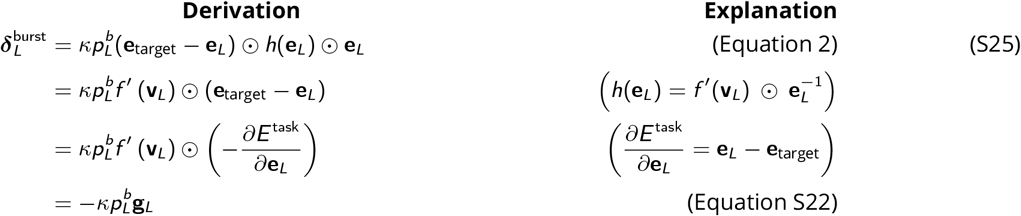

where 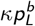 acts as a proportionality constant. For analytical simplicity, we assume throughout this section that *κ* = 1, and the baseline burst probability is homogeneous across neurons within each area, setting 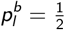. However, these derivations extend to heterogeneous baseline burst probabilities, which we do not consider explicitly here. Using a similar recursive approach to Equation S23 and the Taylor series expansion of the sigmoid function 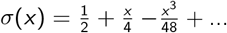, the hidden area burst rate change can be written as:

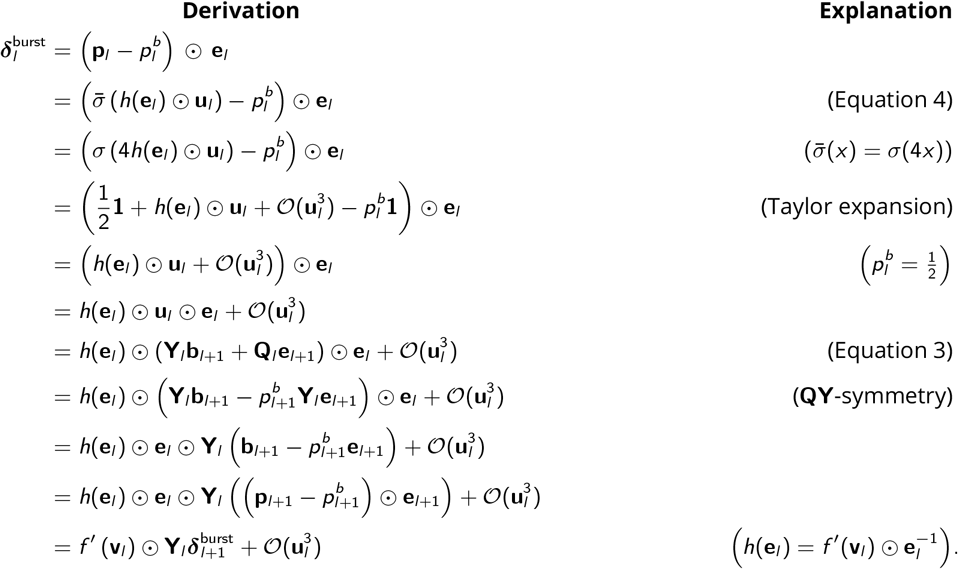

where the *O*(**u**^3^) term represents the approximation error arising from the higher-order remainder of the Taylor expansion. In general, the remainder is cubic in the argument of the dendritic nonlinearity, *h*(**e**_*l*_ ) ⊙ **u**_*l*_, and therefore depends on **u**_*l*_ as well as the scaling induced by *h*(**e**_*l*_ ), with an additional multiplicative scaling by **e**_*l*_ . For analytical simplicity, we assume that both **e**_*l*_ and *h*(**e**_*l*_ ) are bounded, such that the approximation error remains of order 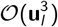. This holds, for example, for sigmoid activations *f* = *σ*, where **e**_*l*_ ∈ [0, 1] and *h*(**e**_*l*_ ) = 1 − **e**_*l*_ is also bounded.

For activation functions such as ReLU, *h*(**e**_*l*_ ) is not bounded in general, and the remainder cannot be reduced to 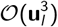 . In this case, the approximation error can be amplified by the inverse event rate term 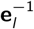 appearing in the definition of *h*, and can become large when event rates are small. Intuitively, this is because error information is multiplexed within the change in burst activity, so low event rates limit the magnitude of gradients that can be represented in the burst rate signal. In practice, this can be mitigated by clipping the inverse-event factor, for example by redefining 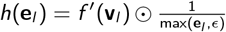, which bounds *h*(**e**_*l*_ ) at the cost of attenuating the propagated error signal at low event rates.

If the apical potentials are sufficiently small, in the sense that ∥**u**_*l*_ ∥_∞_ ≪ 1, then the cubic and higher-order terms 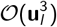 become negligible. If we further assume that the feedback weights are symmetric with the feedforward weights, 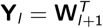, a condition referred to as the **WY**-symmetric configuration, we obtain:

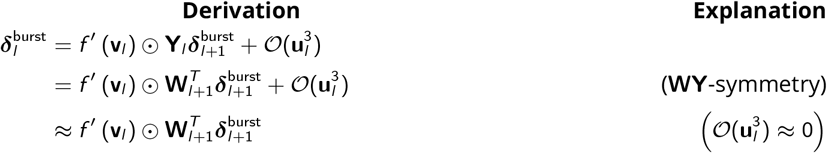

This recursive relationship has the same form as the backpropagation recursion in Equation S23. Thus, using Equation S25 as the base case, it follows by induction for every hidden area that 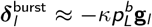. Similarly to backprop in Equation S24, this can be distributed to the weights using 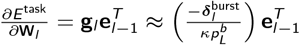. The link between the update rules of BurstCCN and backprop can be seen clearly:

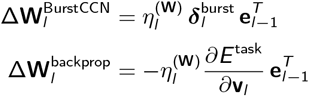

where the learning rate 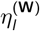 of the BurstCCN absorbs the constant scalar 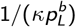. If the **WY**-symmetry assumption is not imposed, the recursive relationship mirrors that of Feedback Alignment (ANN-FA), and the BurstCCN updates approximate the corresponding ANN-FA updates.

